# Synaptotagmin 7 is enriched at the plasma membrane through γ-secretase processing to promote vesicle docking and control synaptic plasticity in mouse hippocampal neurons

**DOI:** 10.1101/2021.02.09.430404

**Authors:** Jason D. Vevea, Grant F. Kusick, Erin Chen, Kevin C. Courtney, Shigeki Watanabe, Edwin R. Chapman

## Abstract

Synaptotagmin (SYT) 7 has emerged as key regulator of presynaptic function, but its localization and precise function in the synaptic vesicle cycle remain unclear. Here, we used iGluSnFR to optically and directly interrogate glutamate release, at the single bouton level, in SYT7 KO dissociated mouse hippocampal neurons. We analyzed asynchronous release, paired pulse facilitation, and synaptic vesicle replenishment, and found that SYT7 contributes to each of these processes to different degrees. ‘Zap-and-freeze’ electron microscopy revealed that loss of SYT7 impairs the docking of synaptic vesicles after a stimulus and the recovery of depleted synaptic vesicles after a stimulus train. To execute these functions, SYT7 must be targeted to the plasma membrane via γ-secretase-mediated cleavage of the amino terminus, followed by palmitoylation. The complex sorting itinerary of SYT7 endows this Ca^2+^-sensor with the ability to control crucial forms of synaptic function and plasticity.

- SYT7 mediated asynchronous release, paired pulse facilitation, and synaptic vesicle replenishment was observed optically at individual hippocampal synapses
- Localization, trafficking, and stability of SYT7 is dependent on processing by γ-secretase
- Short term plasticity defects arise in SYT7KOs due to decreased docking of synaptic vesicles after stimulation
- SYT7 promotes paired-pulse facilitation and asynchronous release via distinct mechanisms

## Introduction

Calcium affords remarkable control over myriad membrane trafficking events in cells. In presynaptic nerve terminals, calcium is particularly important as it regulates numerous aspects of the synaptic vesicle (SV) cycle, including modes of exocytosis, endocytosis, and several forms of synaptic plasticity. There are three modes of exocytosis: synchronous release occurs with a short delay following a stimulus, asynchronous release is characterized by a longer, variable delay following a stimulus, and spontaneous release occurs in absence of electrical activity. The magnitude or rate of these modes can be influenced by previous synaptic activity to mediate various forms of short-term synaptic plasticity (**Barrett & Stevens, 1972**). Calcium is central to regulating SV release (**Katz & Miledi, 1965**) and as such, calcium sensing proteins must govern the SV cycle. The synaptotagmins (SYT) are a family of proteins characterized by the presence of tandem C2 domains that often mediate binding to calcium and phospholipid bilayers (**Wolfes & Dean, 2020**). The most studied isoform is synaptotagmin 1 (SYT1), which promotes rapid synchronous SV exocytosis (**Littleton, Stern, Schulze, Perin, & Bellen, 1993; Geppert et al., 1994**) and clamps spontaneous release (**Littleton et al., 1993; Liu, Bai, Xue, et al., 2014**). SYT2 is a closely related isoform that is expressed in neurons in the cerebellum and spinal cord where it functions in the same manner as SYT1 (**Pang et al., 2006**). Other SYT isoforms are expressed throughout the brain and have distinct affinities for calcium and membranes. Some isoforms do not bind calcium at all while the others fall into three distinct kinetic groupings based on how fast they bind or unbind to membranes in response to changes in calcium [Ca^2+^] (**Hui et al., 2005**).

Recently, synaptotagmin 7 (SYT7) has garnered particular interest because it is now implicated in multiple modes of SV release and at least two forms of synaptic plasticity (**Huson & Regehr, 2020**). SYT7 is widely expressed throughout the body and highly expressed in brain (**C. Li et al., 1995; Sugita et al., 2001**). Interestingly, SYT7 has been reported to be localized to a number of distinct compartments in cells, including the plasma membrane of PC12 cells, presynaptic terminals in neurons (**Sugita et al., 2001**), lysosomes in NRK fibroblasts (**Martinez et al., 2000**), endolysosomal compartments in PC12 cells and neurons (**Monterrat, Grise, Benassy, Hemar, & Lang, 2007**), and dense core vesicles (DCVs) in chromaffin cells (**Fukuda, Kanno, Satoh, Saegusa, & Yamamoto, 2004**). When SYT7KO mice were first generated, they showed grossly normal brain structure and no observable neurological phenotype (**Chakrabarti et al., 2003**). However, inhibition of SYT7 revealed defects in plasma membrane repair (**Reddy, Caler, & Andrews, 2001**), and the first SYT7 knockout (KO) studies found reduced rates of neurite outgrowth (**Arantes & Andrews, 2006**), and alterations in bone density homeostasis (**Zhao et al., 2008**), all stemming from deficiencies in lysosomal exocytosis. Additional studies revealed that changes in SYT7 expression alter DCV exocytosis in PC12 (**Wang, Chicka, Bhalla, Richards, & Chapman, 2005**), adrenal chromaffin (**Schonn, Maximov, Lao, Sudhof, & Sorensen, 2008; Rao et al., 2014**) and pancreatic beta cells (**Gut et al., 2001; Y. Li et al., 2007; Gauthier et al., 2008; Gustavsson et al., 2008**).

Early experiments, in which SYT7 was over-expressed (OE) in neurons hinted at a role for SYT7 in the SV cycle by uncovering a complex endocytosis phenotype (**Virmani, Han, Liu, Sudhof, & Kavalali, 2003**). However, subsequent electrophysiological examination of synaptic transmission concluded that there was no change in SV release or short-term synaptic plasticity in the SYT7KOs (**Maximov et al., 2008**). This was unexpected, because the high affinity of SYT7 for calcium and its slow intrinsic kinetics made this isoform a compelling candidate to serve as a calcium sensor for asynchronous release or for short-term plasticity (**Hui et al., 2005**). Consequently, in 2010, a role for SYT7 in asynchronous release during high frequency train stimulation (HFS) was described at the zebrafish neuromuscular junction (**Wen et al., 2010**) and then in hippocampal neurons from mice (**Bacaj et al., 2013**). Based on these studies, asynchronous release appears to be mechanistically distinct when it occurs following a single stimulus versus a stimulus train, again because loss of SYT7 appears to impact release only when more than one stimulus is given. Interestingly, SYT7 has been shown to promote asynchronous release from neurons after a single stimulus, but only after artificial, ectopic expression of SNAP-23 (**Weber, Toft-Bertelsen, Mohrmann, Delgado-Martinez, & Sorensen, 2014**). At the same time, SYT7 was found to mediate calcium dependent SV replenishment in response to HFS (**Liu, Bai, Hui, et al., 2014**). The Regehr group then discovered that SYT7 is required for paired-pulse facilitation (PPF) in Schaffer collateral, corticothalmic, mossy fiber, and perforant pathway synapses (**Jackman, Turecek, Belinsky, & Regehr, 2016**). These authors also found that facilitation supported frequency invariant transmission at Purkinje cell and vestibular synapses (**Turecek, Jackman, & Regehr, 2017**). At granule cell synapses, they also observed a role for SYT7 in facilitation and asynchronous release (**Turecek & Regehr, 2018**). Finally, a role for SYT7 in facilitation, asynchronous release, and SV replenishment was observed at GABAergic basket cell-Purkinje cell synapses (**Chen, Satterfield, Young, & Jonas, 2017**).

The function of SYT7 has gone from no identified role in the SV cycle, to fulfilling several different roles that likely involve the regulation of similar processes at many kinds of synapses. To reconcile these phenotypes, and to gain insight into the underlying mechanisms, we examined SV exocytosis in wild-type and SYT7KO hippocampal synapses in dissociated cultures using an optical biosensor for glutamate (iGluSnFR) (**Marvin et al., 2018**). Moreover, to gain insights into precise steps in the SV cycle that are regulated by SYT7, we carried out ‘zap-and-freeze’ (**Kusick et al., 2020**) electron microscopy experiments. Use of iGluSnFR allowed us to monitor glutamate release directly from single presynaptic nerve terminals, and ‘zap-and-freeze’ EM yielded novel insights into the membrane trafficking events that occur within 5 ms of an action potential. Furthermore, we examined the localization, post-translational modifications, and function of SYT7 in neurons using powerful new JaneliaFluor (**Grimm et al., 2017**) HaloTag ligands in conjunction with SYT7 retargeting strategies. We show that synapses lacking SYT7 exhibit subtle defects in asynchronous release, a complete disruption of PPF, and decreased rates of SV replenishment. We propose that these deficiencies originate, at least in part, from severe reductions in SV docking during activity, as revealed by ‘zap-and-freeze’ electron microscopy in which images are captured within milliseconds after a of stimulus. Surprisingly, we discovered that the amino terminus of SYT7 is cleaved by the Alzheimer’s disease relevant γ-secretase complex; the stability and localization of SYT7 is dependent on this proteolytic processing step and concurrent palmitoylation. We propose that these modifications are critical for the subsynaptic membrane trafficking of SYT7 and its role in supporting the SV cycle. Finally, by retargeting and restricting SYT7 to various membranes in the synapse, we show for the first time that SYT7 must localize to the plasma membrane (PM) to support asynchronous release, PPF, and SV replenishment.

## Results

### SYT7 influences presynaptic neurotransmitter release during short term synaptic plasticity

To monitor SV exocytosis, we transduced the low affinity (S72A) optical glutamate reporter iGluSnFR (**Marvin et al., 2018**) into cultured mouse hippocampal neurons. This allowed us to monitor glutamate release irrespective of confounding postsynaptic factors (**Wu et al., 2017**). Using a single stimulus, we analyzed and compared the magnitude of glutamate release between wild type (WT) and SYT7KO neurons, as well as the balance of synchronous and asynchronous release. No differences in the magnitude of glutamate release between WT and SYT7KO neurons were observed. Representative traces are shown in (Fig. 1a) with peak ΔF/F_0_ quantitation in (Fig. 1b). We used a 10 ms cutoff to distinguish between synchronous and asynchronous glutamate peaks, as described in earlier patch clamp experiments (**Yoshihara & Littleton, 2002; Nishiki & Augustine, 2004**). We found a small (3% difference in medians or 1.8% according to the Hodges-Lehmann estimate), but statistically significant decrease, in asynchronous release from SYT7KO neurons in response to a single stimulus (Fig 1c). Previous comparisons examining release, triggered by a single action potential and monitored electrophysiologically, found no differences between WT and SYT7KO synapses (**Liu, Bai, Hui, et al., 2014; Chen et al., 2017**). The small change that we detected is likely due to the sensitivity afforded by using the iGluSnFR optical probe to directly monitor glutamate release, as compared to post synaptic recordings.

**Figure 1:**
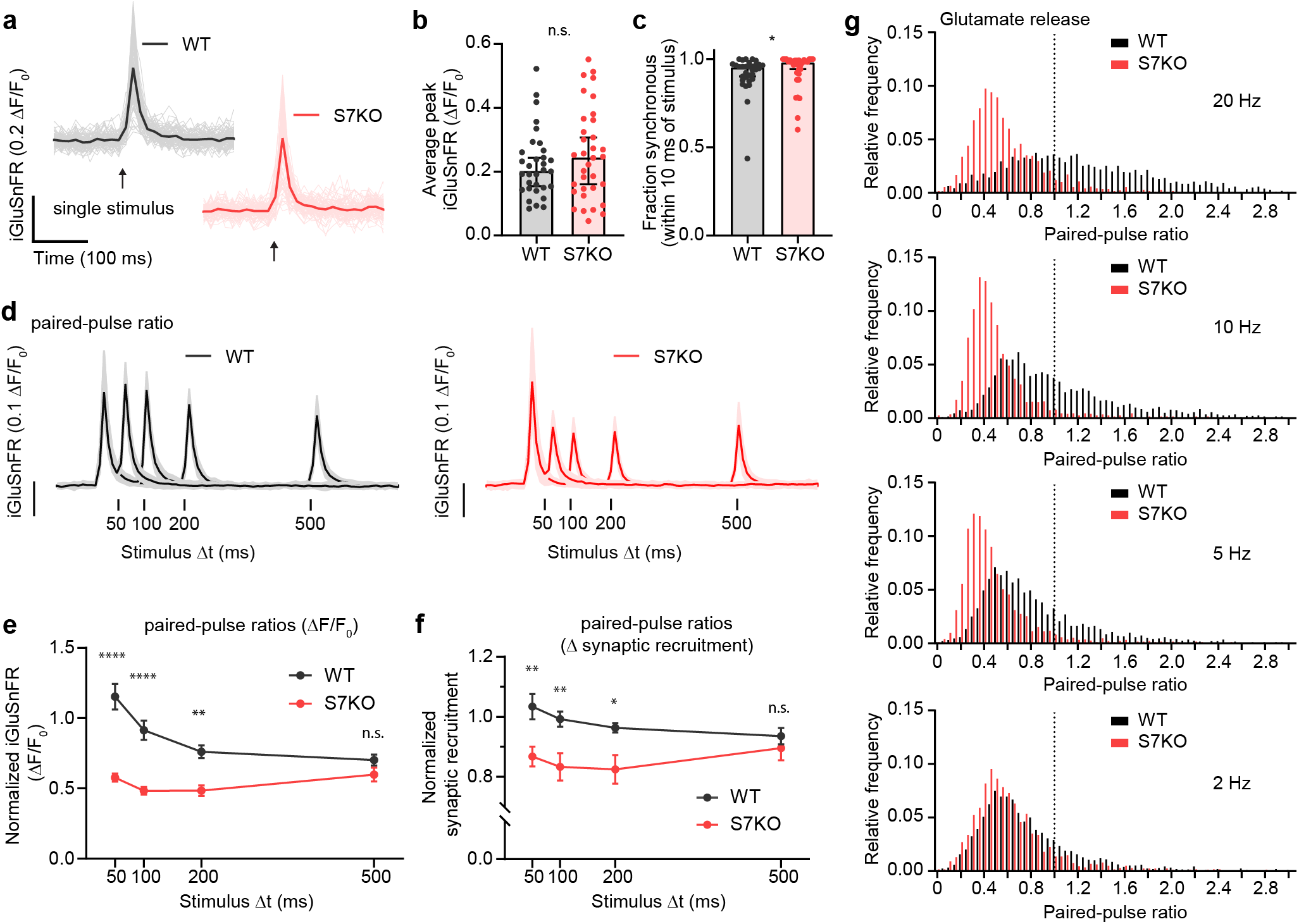
SYT7 influences presynaptic neurotransmitter release during short term synaptic plasticity. **a**) Representative sf iGluSnFR S72A (hereon iGluSnFR) traces from single stimulus experiments. Lighter traces are individual regions of interest (ROIs) and dark, bold traces are the average of all light traces from a full field of view (FOV), single stimulus denoted with arrow. WT in black and grey and SYT7KO in red and light red. **b**) Peak iGluSnFR signals between WT (0.203 [95% CI 0.154-0.244] ΔF/F_0_) and SYT7KO (0.245 [95% CI 0.160-0.308] ΔF/F_0_). Values are medians with 95% CI representing error, Mann-Whitney test p = 0.4554, each n is a separate FOV (n = 32 (WT) and 34 (SYT7KO) from 4 independent experiments). **c**) Fraction of synchronous release, defined as peak iGluSnFR signals arriving within 10 ms of stimulus from total release of 500 ms following stimulus, compared between WT (0.9522 [95 % CI 0.902-0.965]) and SYT7KO (0.9808 [95% CI 0.943-0.993]). Data from same n as in b). Values are medians with 95% CI representing error, Mann-Whitney test p = 0.0326. **d**) Average traces from paired-pulse ratio (PPR) experiments with four interstimulus intervals compared. WT in black with grey standard deviation as error and SYT7KO in red with light red standard deviation as error, n = 14 (WT 20 Hz), 14 (WT 10 Hz), 15 (WT 5 Hz), 13 (WT 2 Hz), 15 (SYT7KO 20 Hz), 13 (SYT7KO 10 Hz), 14 (SYT7KO 5 Hz), 13 (SYT7KO 2 Hz) from 3 independent experiments. **e**) Quantification of PPR (peak iGluSnFR ΔF/F_0_) from WT (black) and SYT7KO (red), values are means +/− SEM. P-values are as follows **** < 0.0001, ** = 0.0012, by two-way ANOVA with Sidak’s multiple comparisons test, full statistics are provided in supplementary statistics table 1. **f**) Quantification of fractional active synapses, i.e. number of synapses demonstrating peak release above baseline during second stimulus of paired pulse. Values are means +/− SEM. P-values are, in order from left to right, ** = 0.0052, ** = 0.0099, * = 0.0289, by two-way ANOVA with Sidak’s multiple comparisons test, full statistics are provided in supplementary statistics table 2. **g**) Relative frequency histograms of paired-pulse ratio (PPR) from all ROIs quantified from PPR trials, 20 Hz, 10 Hz, 5 Hz, 2 Hz, WT (black) and SYT7KO (red). Vertical dotted line delineates a PPR of 1.

Next, we examined a form of short-term synaptic plasticity. In a variety of synapses, for a brief period of time after a conditioning first stimulus, release is enhanced if a second stimulus is applied, and this is termed paired-pulse facilitation (PPF) (**Regehr, 2012**). We note that the ratio of the first two responses is more generally termed the paired-pulse ratio (PPR). Here, we examined the PPF tuning window by interrogating glutamate release at 50, 100, 200, and 500 ms interstimulus intervals. For WT synapses, we detected facilitation (~110%) using iGluSnFR at 50 ms interstimulus intervals, a mild decline at 100 ms, and loss of PPF at 200 and 500 ms interstimulus intervals (Fig. 1d). In SYT7KO neurons, PPF is absent (Fig. 1d); hence this simplified system recapitulates the role of SYT7 in PPF that was reported using hippocampal slice preparations (**Jackman et al., 2016**). Quantifying the PPR, we found that SYT7KOs release approximately half the amount of glutamate in response to the second stimulus relative to the first stimulus at all intervals (Fig. 1e). As emphasized above, no differences were observed when quantifying the magnitude of glutamate release triggered by the first stimulus between WT and SYT7KO neurons; again, differences emerged only after the second stimulus (Fig. S1a-b). An advantage of the optical measurements utilized here is that they report the spatial distribution of transmission and can report the number of active synapses from one response to the next (synaptic recruitment). Interestingly, in WT neurons, the number of synapses that actively release glutamate in response to a conditioning pulse are maintained, while SYT7KO neurons deactivate ~10% of synapses following interstimulus intervals of 50, 100, and 200 ms. Release in the WT and SYT7KO become equal only at the 500 ms interstimulus interval (Fig. 1f, Fig. S1c-f). Visualizing 20 Hz PPF (Fig. S1g-h), it is readily evident that there is a near universal decrease in the ability of SYT7KO synapses to release glutamate following a conditioning stimulus. A temporally color-coded max projection of the first two stimuli from a 20 Hz pulse (50 ms interval) is shown in Fig. S1g. Release triggered by the first stimulus is color coded green and release from the second stimulus is color coded magenta. Facilitation is visible as white or magenta while depression is visible as green. The relative frequency distribution of paired pulse ratios for 50, 100, 200 and 500 ms interstimulus intervals are shown in (Fig. 1g) where facilitating (PPR > 1) and depressing (PPR < 1) synapses are readily observable. These findings demonstrate that PPF is directly mediated by an enhancement of glutamate release from already active presynaptic boutons and not via recruitment of previously silent boutons. Hence, in WT synapses, SYT7 must somehow promote SV fusion during activity, perhaps by enhancing docking or stabilizing a docking intermediate; we address these possibilities further below.

### SYT7 counteracts synaptic depression and promotes asynchronous release during sustained stimulation

We further examined glutamate release from hippocampal synapses using iGluSnFR as described in Fig. 1, but now as a function of high frequency stimulus trains (HFS). The HFS consisted of 50 stimuli at 20 Hz (2.5 s stimulation epoch), which was sufficient to reach steady state depression. Representative traces from individual ROIs are shown in (Fig. 2a) from WT (i) and SYT7KO (ii) neurons. Average iGluSnFR traces comparing WT and SYT7KO neurons during HFS show broad depression and loss of tonic charge at SYT7KO synapses (Fig. 2b). This was similar to findings obtained via electrophysiological recordings of EPSCs (**Liu, Bai, Hui, et al., 2014**). During HFS, average glutamate release declines in WT neurons but this depression occurs more rapidly and deeply in SYT7KO neurons (Fig. S2a) (**Liu, Bai, Hui, et al., 2014**). Similarly, the number of active SYT7KO synapses that release glutamate also declined significantly (Fig. 2c and S2a). By measuring the cumulative iGluSnFR signal during a HFS, we calculated the SV replenishment rate (Fig. 2d). This rate is the slope of a linear regression fitted to steady state that is reached during the last 1.5 s of the HFS. SYT7KO synapses replenish SVs at about half the rate of WT synapses (Fig.2e), similar to what has been reported previously from electrophysiological measurements (**Liu, Bai, Hui, et al., 2014; Chen et al., 2017**). Interestingly, from the single ROI traces shown in Fig. 2a, once release reaches steady state, fluorescent iGluSnFR responses decayed to baseline before the next stimulus. This contrasts to the average iGluSnFR fluorescence change which does not decay to baseline (Fig. 2b). We argue that the failure of the signal to decay to baseline in our average traces is analogous to the tonic charge component measured via electrophysiology and represents asynchronous release from single synapses, that when averaged, forms the tonic charge component. Therefore, we can use iGluSnFR imaging to monitor individual ROIs and measure HFS related asynchronous release. Indeed, a smaller amount of asynchronous release is triggered during HFS at SYT7KO synapses relative to WT synapses, and this difference widens as stimulation progresses (Fig. 2f).

**Figure 2:**
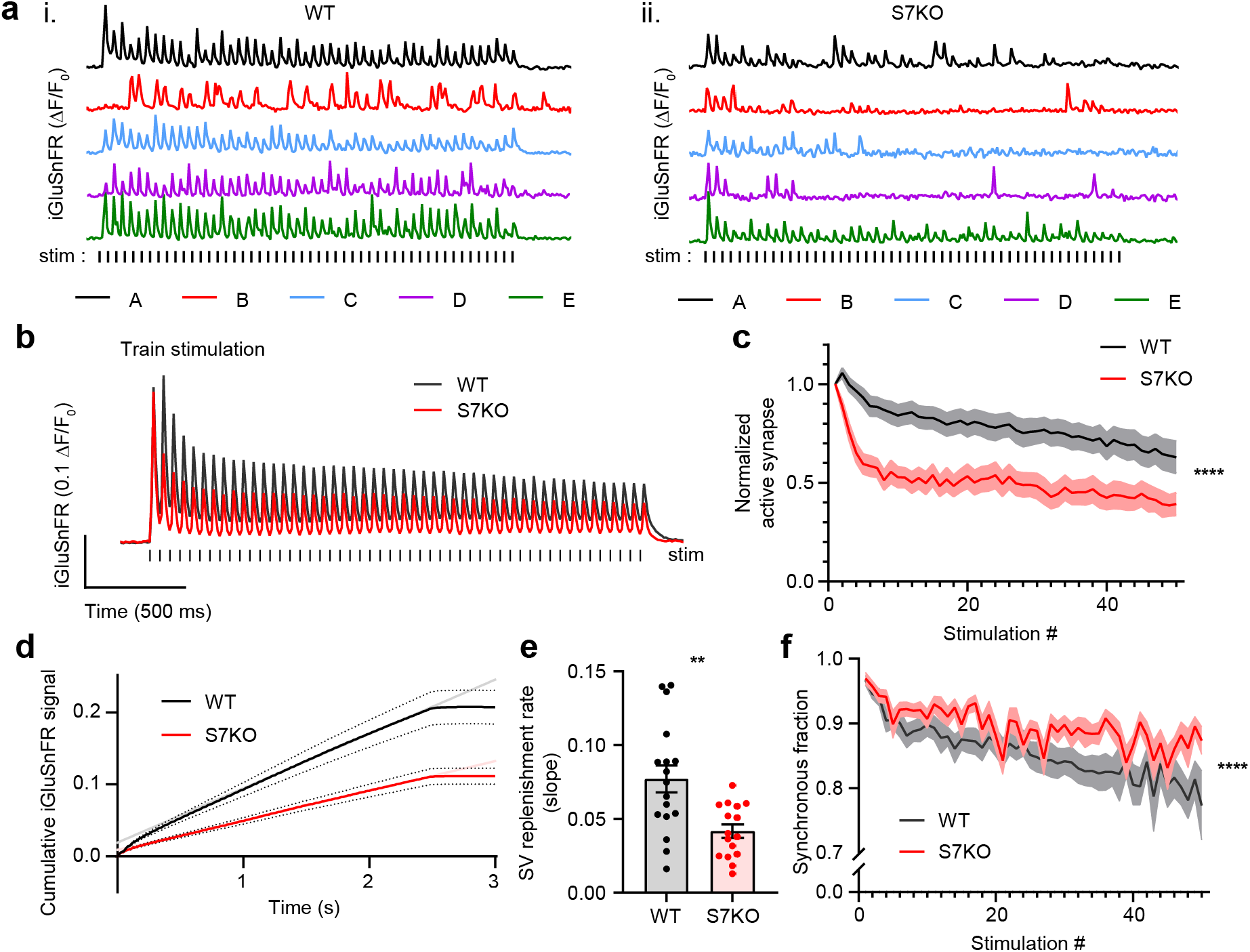
SYT7 counteracts depression and promotes asynchronous release during sustained stimulation. **a**) Representative traces of iGluSnFR ΔF/F_0_ signals (single ROIs A-E), from one FOV during high frequency stimulation (HFS) of WT (i) and SYT7KO (ii) neuronal preparations. Samples were field stimulated with a frequency of 20 Hz for 2.5 s (50 APs). **b**) Average iGluSnFR ΔF/F_0_ traces during HFS for WT (black, n = 17) and SYT7KO (red, n = 16), from 3 independent experiments (same source data for Fig. 2b – 2f). **c**) Fraction of active synapses, defined as synapses releasing peak glutamate above baseline, >4 SD above noise, as a function of stimulation number during HFS. Values are means (lines) +/− SEM (lighter shade error), p < 0.0001 by two-way ANOVA comparing genotype. **d**) Plot of average cumulative iGluSnFR ΔF/F_0_ signal from WT (black) and SYT7KO (red) neurons vs time. Dotted lines represent SEM and grey (WT) and light red (SYT7KO) linear lines represent linear fits to the last 1.5 seconds of the train. **e**) SV replenishment rates were calculated from slopes of linear regressions from individual traces used in d). Values are means +/− SEM, WT (0.077 +/− 0.009) and SYT7KO (0.042 +/− 0.004), ** = 0.0019 p-value using unpaired, two-tailed t-test. **f**) Fraction of synchronous release, defined as peak iGluSnFR ΔF/F_0_ within 10 ms of stimulus from a total of 50 ms post stimulus (i.e. interstimulus interval) as a function of stimulation number during HFS. Values are means (bold lines) +/− SEM (lighter shade fill), p < 0.0001 by two-way ANOVA comparing genotype.

### SYT7 helps maintain docked and total synaptic vesicle pools after stimulation

To directly visualize the events that occur at synapses during HFS, and to understand how SYT7KO synapses depress faster, have less asynchronous release, and exhibit a much slower SV replenishment rate, we turned to ‘zap-and-freeze’ electron microscopy (EM). This technique involves freezing synapses as fast as 5 milliseconds after electrical stimulation, followed by freeze substitution and EM to observe synaptic ultrastructure. At rest, SYT7KO synapses have no gross morphological defects, with a normal complement of docked (in contact with the active zone plasma membrane) and total SVs in boutons (Fig. 3a-c). In WT synapses, 40% of docked vesicles are lost after a single action potential, as previously reported (**Kusick et al., 2020**). The number of docked vesicles remains similar 5 ms after HFS, presumably because docked vesicle recovery matches depletion. By 5 s after HFS, the number of docked vesicles partly recover to baseline. A similar sequence of loss and recovery of docked vesicles was observed in SYT7KO synapses. However, in all conditions after stimulation, SYT7KO synapses had 30-40% fewer docked vesicles than the corresponding condition in WT (Fig. 3a-c). It should be noted that this increased loss of docked vesicles is not due to increased depletion of vesicles by exocytosis, as indicated by iGluSnFR measurements above (Fig. 2).

**Figure 3:**
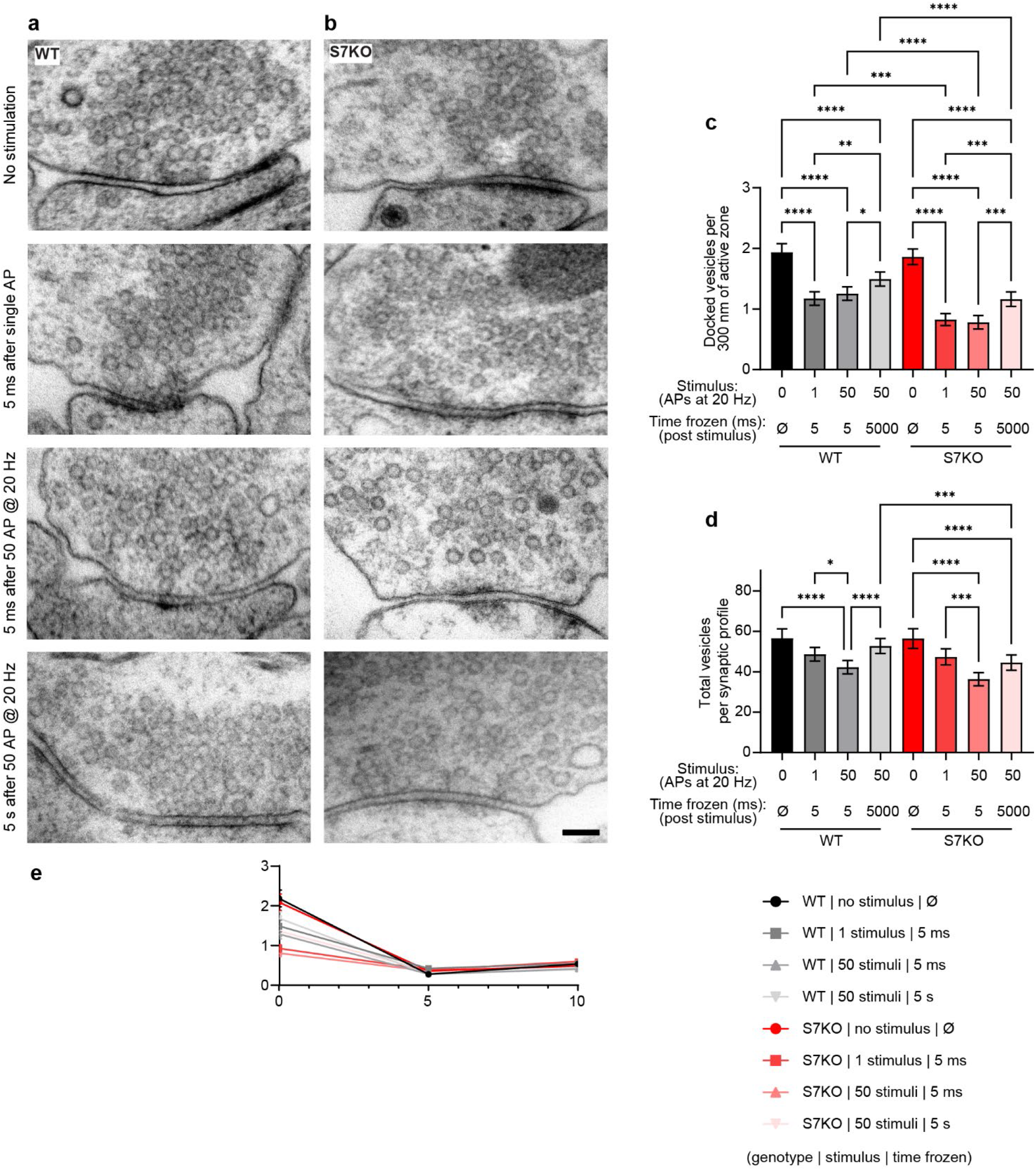
SYT7 enhances synaptic vesicle docking after stimulation. Representative electron micrographs of high-pressure frozen **a**) WT and **b**) SYT7KO synapses from labeled conditions. Scale bar = 100 nm. **c**) Quantification of docked vesicle number normalized to 300 nm of active zone at rest, after stimulation with 1 AP, or 50 APs, and then frozen at 5 ms post-stimulus or 5 s post-stimulus. Docked vesicles are defined in high-pressure frozen samples as in contact with the plasma membrane at the active zone (0 nm between the plasma membrane and vesicle membrane). WT conditions are in black to grey and SYT7KO conditions are red to pink. Values are means +/− 95% CI and are from 3 biological replicates and over 300 n per condition (n = individual 2D EM images). All comparisons and summary statistics are provided in supplementary statistics table 4, p-values are as follows **** < 0.0001, *** < 0.001, ** < 0.01, * = 0.05, by Kruskal-Wallis test with Dunn’s multiple comparison correction. **d**) Quantification of vesicle number per 2D synaptic profile at rest, after stimulation with 1 AP, or 50 APs, and then frozen at 5 ms post-stimulus or 5 s post-stimulus. Values are means +/− 95% CI and are from 3 biological replicates and over 2500 individual 2D EM images. All comparisons and summary statistics are provided in supplementary statistics table 5, p-values are as follows **** < 0.0001, *** < 0.001, ** < 0.01, * = 0.05, by Kruskal-Wallis test with Dunn’s multiple comparison correction. **e**) Quantification of SV number in relation to distance from active zone (a.z.) up to 100 nm. Inset denoted with # sign is enlarged to show SV distribution in close proximity to a.z. Values are means +/− SEM.

In response to a single stimulus, WT and SYT7KO neurons do not display any decreases in total SV number. However, following HFS, at the 5 ms time point, a modest decrease was observed in both conditions, and while WT synapses recovered 5 s after HFS, SYT7KO synapses did not (Fig. 3d). Importantly, careful analysis of the distribution of SVs within 100 nm of the active zone revealed that there were no changes, other than the docked pool, under any condition in WT or SYT7KO synapses (Fig. 3e). This result demonstrates the reduction in docking is specific and is not secondary to the reduction in the total number of vesicles near active zones. The comparison of WT and SYT7KO synapses by ‘zap-and-freeze” revealed two important observations that may help explain the complicated synaptic phenotype of the KO. SYT7KO synapses display a greater loss of docked vesicles after a single stimulus and after HFS. Docking is a prerequisite to fusion, so decreases in docked vesicles after a stimulus could account for decreased asynchronous release, decreased PPF, and increased depression during HFS. Additionally, compared to WT synapses, SYT7KO synapses exhibit a decrease in the total number of SVs 5 s after HFS. This suggests that not only do SYT7KO synapses display a docking defect, but also suffer from a SV reformation defect lasting seconds after an HFS. SV docking and SV reformation are presumably two different processes and take place in different areas of the presynapse. To understand how SYT7 influences both processes it is crucial to characterize the localization and trafficking of this protein.

### In hippocampal neurons, SYT7 is localized to both the axonal plasma membrane and LAMP1+ organelles that include active lysosomes

As outlined in the Introduction, SYT7 localizes to several distinct subcellular compartments in a variety of cell types; its exact localization in mature neurons remains unclear. In dissociated hippocampal neurons, endogenous SYT7 is not detectable above background fluorescence by immunocytochemistry (ICC), presumably resulting from a mix of low expression and poor antibody performance. So, we over-expressed, untagged SYT7α, via sparse lentiviral transduction, and found that it localized to axons and LAMP1+ structures (Fig. 4a). Importantly, we used low levels of lentivirus so that we overexpressed just enough protein to detect with the antibody. The location of a carboxy labeled SYT7α-HaloTag, with respect to a cytosolic fluorescent marker (Fig. 4b) and a plasma membrane targeted msGFP (Fig. S3a), demonstrates the clear polarized distribution of SYT7 to axons but not dendrites; this also confirms that a carboxy-HaloTag does not interfere with SYT7 trafficking. Axonal plasma membrane localization was further established via super resolution Airyscan imaging. Here, SYT7α-HaloTag is resolved from the axoplasma (Fig. 4d-e and S3b-c). The line profile reveals cytosolic-mRuby3 signal peaking in the middle of two characteristic ‘double bump’ signals from SYT7α-HaloTag, which resides on the plasma membrane. Because SYT7α is localized to the axon and influences the SV cycle, it is reasonable to predict that SYT7α may be translationally regulated, akin to bona fide SV proteins. The translation of SV proteins is correlated with synaptogenesis, so we probed synaptophysin (SYP), synaptotagmin 1 (SYT1), SYT7, and total protein as a function of development and found that while SYP and SYT1 protein levels rise together, SYT7 does not follow the same trend as these SV proteins (Fig. S3d-e). Additionally, we expressed SYT7α-HaloTag in HEK293T cells along with cytosolic msGFP and LAMP1-mRuby3. We found that SYT7α-HaloTag localized to the plasma membrane and lysosomal structures in these non-neuronal cells (Fig. S3f).

**Figure 4:**
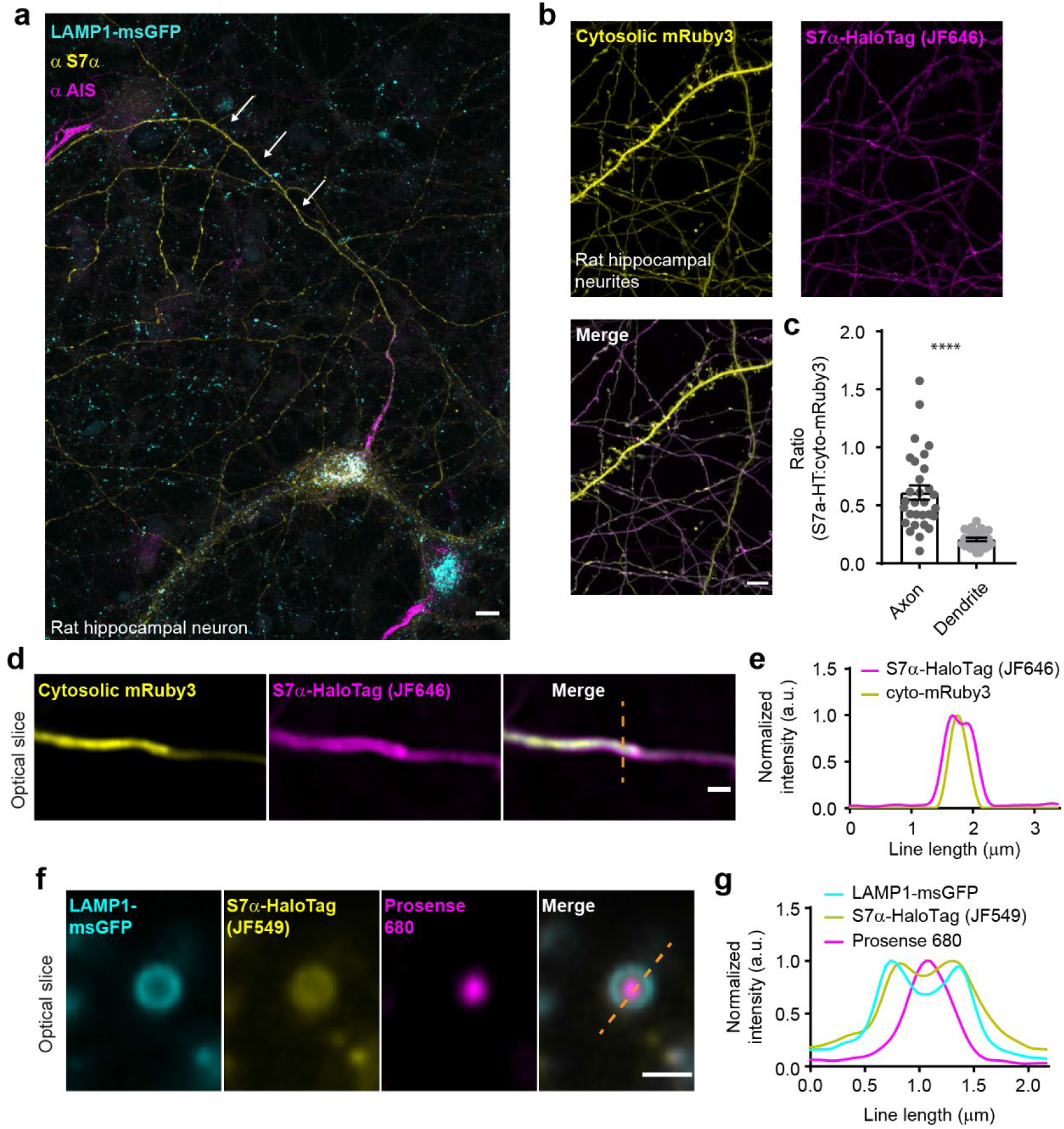
In hippocampal neurons, SYT7 is localized to both the axonal plasma membrane and LAMP1+ organelles that include active lysosomes. **a**) Representative super-resolution fluorescent immunocytochemistry (ICC) image of rat hippocampal neurons at 15 DIV expressing uniformly transduced LAMP1-msGFP and sparsely transduced, untagged, SYT7α. These neurons were fixed and stained with antibodies to SYT7 (juxta-membrane region), and the axon initial segment (AIS) (anti pan-neurofascin). Scale bar 5 microns. **b**) Representative super-resolution images of cytosolically expressed mRuby3 (yellow/top left), Syt7α-HaloTag/JF646 (magenta/top right), and merged (bottom left). Scale bar 5 microns. **c**) Quantification of the ratio between fluorescent channels. Axonal ratio of SYT7-HaloTag:mRuby3 signal is 0.61 +/− 0.06, n = 30, while dendritic ratio is 0.21 +/− 0.01. Values are means +/− SEM from 2 independent experiments, p-value < 0.0001 using unpaired, two tail, Welch’s t-test. **d**) Representative super-resolution optical slice of an axon (identified via morphology) expressing cytosolic mRuby3 (yellow) and SYT7α-HaloTag/JF646 (magenta). Merged image also denotes line used in e). **e**) Graph plotting the normalized intensity profile along the orange dashed line in d). Scale bar 1 micron. **f**) Representative super-resolution optical slice of a somatic lysosome from a rat hippocampal neuron at 16 DIV expressing LAMP1-msGFP (cyan), SYT7α-HaloTag/JF549 (yellow), and incubated with 0.5 μM Prosense 680 (magenta) for 12 hours. Merged image also denotes line used in g). **g**) Graph plotting the normalized intensity profile along the dashed orange line in f). Scale bar 1 micron.

To examine SYT7 localization in mature neurons with respect to the endo-lysosomal system, we co-expressed SYT7α-HaloTag and LAMP1-msGFP. Indeed, we observed broad colocalization of these constructs (Fig. S3g-i). LAMP1-msGFP identifies mature lysosomes as well as intermediates in the endo-lysosomal compartment (**Cheng et al., 2018**). To specifically identify active lysosomes, we incubated neurons with Prosense 680. This molecule is self-quenching and membrane impermeant; when cleaved by lysosomal proteases it dequenches, and thus fluorescently labels active lysosomes (**Weissleder, Tung, Mahmood, & Bogdanov, 1999**). Interestingly SYT7α-HaloTag was present throughout the endo-lysosomal compartments, on active and inactive lysosomes (Fig. 4f-g). Importantly, SYT7α-HaloTag is clearly limited to the lysosomal membrane and does not appear to simply colocalize with lysosomes via a degradation pathway.

SYT7 has been localized to lysosomes in non-neuronal cells where it plays a role in lysosomal exocytosis (**Martinez et al., 2000**) and in trafficking cholesterol to the plasma membrane (**Chu et al., 2015**). Cholesterol is a lipid that is critically important for the formation of SVs; cholesterol also binds and regulates interactions between some SV protein (**Thiele, Hannah, Fahrenholz, & Huttner, 2000**). This link between SYT7 function, cholesterol trafficking, and the SV cycle is attractive because it might explain some of the SV cycle related phenotypes of SYT7 null neurons. Knowing that SYT7 resides on the lysosomal membrane in neurons, we investigated cholesterol levels, as well as interactions that are sensitive to changes in the abundance of this lipid, in SYT7KO neurons. More specifically, the SV proteins SYP and synaptobrevin (SYB) interact in a cholesterol dependent manner (**Mitter et al., 2003**). If loss of SYT7 results in decreased trafficking of cholesterol to the plasma membrane as it does in HEK293T cells (**Chu et al., 2015**), we should observe decreased interactions. Using mature neurons and a chemical crosslinker previously shown to successfully probe SYP/SYB interactions (**Mitter et al., 2003**), we did not observe decreased interactions in SYT7KO neurons relative to WT (Fig. S4a-c). Similarly, we do not see a change in any lipid species by thin layer chromatography (Fig. S4d-e), or a buildup of neutral lipids in lysosomes (Fig. S4f). Based on these data, we conclude that SYT7 likely influences the SV cycle from its location on the axonal plasma membrane, and not indirectly by altering the abundance or distribution of cholesterol in neurons. How SYT7 becomes enriched in axons, and how it persists on the axonal plasma membrane in spite of robust membrane cycling during exo- and endocytosis, are questions that we explore in the next series of experiments.

### Synaptotagmin 7 is cleaved by the intramembrane aspartyl protease presenilin

In our efforts to localize SYT7α, we transduced neurons with a variety of tags at its amino- and carboxy-termini. When examining the expression levels of these constructs by immunoblot analysis, we observed that the amino terminally tagged constructs existed as a mix of proteins with the predicted (large) molecular weight of the fusion protein along with bands of apparently the same molecular weight of the untagged protein. The carboxy terminal tagged constructs yielded a single band that corresponded to the size of the full-length protein plus the tag (Fig. S5a-b). Therefore, the artificial N-terminal tag is cleaved off by a cellular protease, and this cleavage must occur near the tag junction, or within the amino terminus of SYT7α. Changing the tag, linker, or deleting the luminal domain, did not affect cleavage of SYT7 (data not shown), thus leaving the transmembrane domain (TMD) as the only possible cleavage site. Cytosolic-side cleavage is unlikely as there are palmitoylation sites on that side of the TMD that influence localization in fibroblasts (**Flannery, Czibener, & Andrews, 2010**).

There are a limited number of intramembrane proteases in cells. Interestingly, with their short luminal tail segments, SYTs have been postulated to be targets of the γ-secretase complex (**Sudhof, 2002**); we tested this idea using inhibitors. Remarkably, a combination of 1 μM DAPT (competitive presenilin inhibitor) and 20 μM GI 254023X (ADAM10 metalloprotease inhibitor) strongly inhibited proteolytic processing of the HaloTag-SYT7α construct (Fig. S5c). DAPT alone appeared to only prevent processing of an already cleaved form of HaloTag-SYT7α; GI 254023X had to be present as well to prevent the cleavage of HaloTag-SYT7α protein. However, when we transduced neurons with untagged SYT7α, only DAPT shifted the mobility of SYT7α on SDS-PAGE (Fig. S5d). These findings indicate that there are two cleavage reactions, one pertaining to the artificial tag, and another targeting the untagged SYT7α protein. Endogenous SYT7 is alternatively spliced to create at least three different isoforms with varying juxtamembrane linker lengths (**Fukuda, Ogata, Saegusa, Kanno, & Mikoshiba, 2002**). We found that DAPT, but not GI 254023X shifted the apparent size of all SYT7 isoforms from rat hippocampal neurons (Fig. 5a). This suggests that SYT7 isoforms do not need to be preprocessed by a metalloprotease and are bona fide direct targets of the γ-secretase complex.

**Figure 5:**
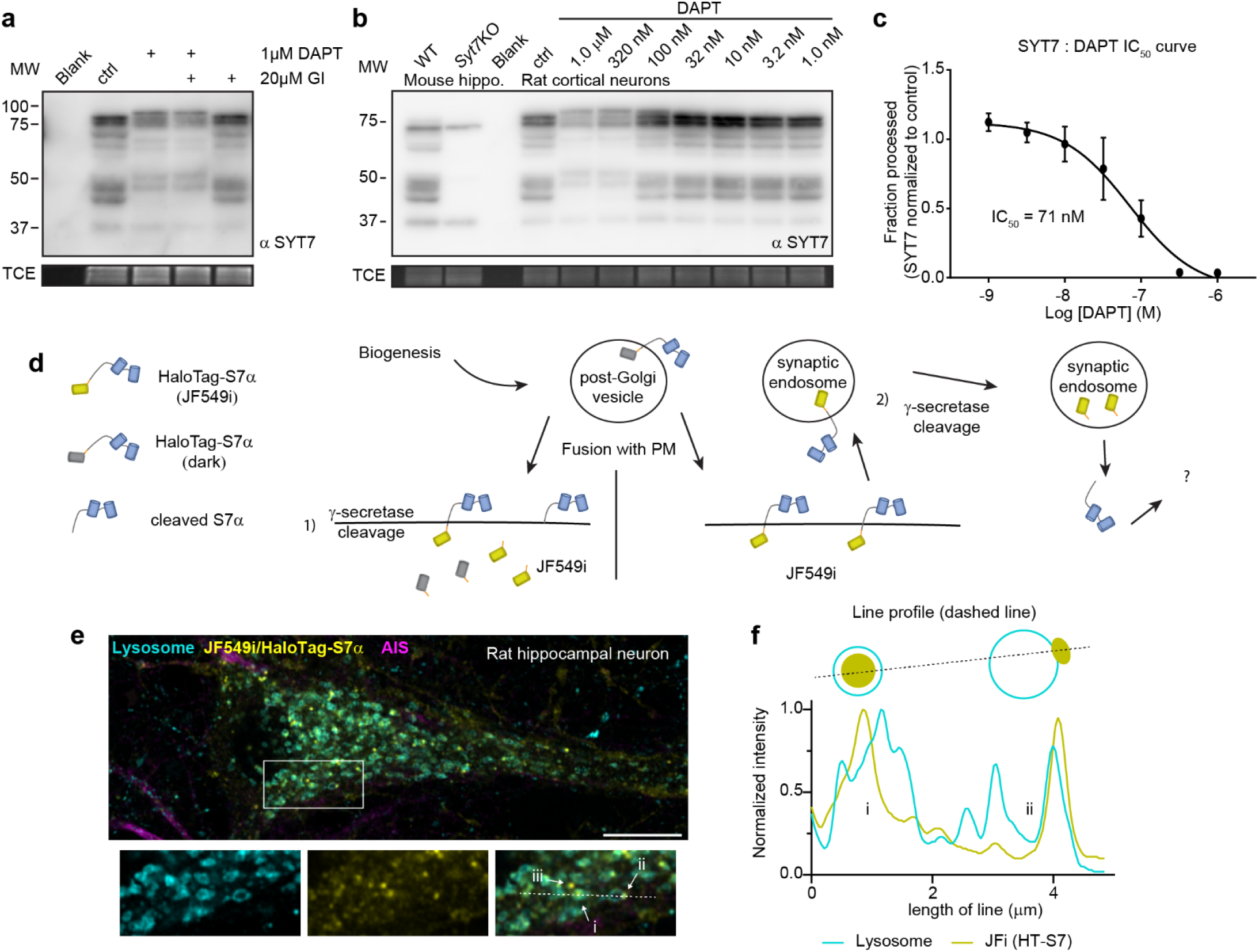
Synaptotagmin 7 is cleaved by the intramembrane aspartyl protease presenilin. **a**) Representative anti-SYT7 immunoblot from rat hippocampal neurons with trichloroethanol (TCE) as a loading control. Conditions from left to right are blank/no protein, control conditions, neurons grown in the presence of 1 μM DAPT (presenilin competitive inhibitor), grown with DAPT and 20 μM GI 254023X (ADAM10 selective inhibitor), and neurons grown in only GI, all from DIV 5 onward. **b**) Representative anti-SYT7 immunoblot using mouse hippocampal neurons for WT and SYT7KO antibody controls along with rat cortical neurons grown in various concentrations of DAPT to assay IC_50_. TCE as a loading control. **c**) Graph of the fractional processed SYT7 when grown in various DAPT concentrations in relation to control conditions (IC_50_ curve) results in an IC_50_ of 71 nM. The lowest specific SYT7 band was used for quantitating cleavage and IC_50_ of DAPT. Values are means +/− SD after log transformation from 3 independent experiments. **d**) Cartoon illustrating logic and methodological approach to determine if full length SYT7 protein transits through the plasma membrane prior to amino terminal cleavage by γ-secretase. JF549i is a membrane impermeant version of JF549 (JF549 and JF549i are nonfluorogenic). In 1., cleavage can take place in the post-Golgi vesicle, prior to axonal plasma membrane (PM) localization or cleavage happens at the PM. No fluorescent HaloTag is observable in this scenario. In 2., SYT7 transits through the PM before being cleaved in a synaptic endosome. Only in this scenario will fluorescent HaloTag be observable in neurons. **e**) Representative super-resolution optical slice of a rat hippocampal neuron transduced with LAMP1-msGFP (cyan) and HaloTag-SYT7α (yellow). Before fixing neurons, they were incubated with 1 nM JF549i for 2 days. Fixed neurons were decorated with anti-pan-neurofascin (magenta) antibodies to mark the AIS. White box indicates area that was enlarged to show detail below the image. The labels i, ii, and iii, indicate areas where JF549i appears inside lysosomes, clustered on the edge of lysosomes, or completely independent of lysosomes, respectively. **f**) Line profile from dashed line in h) with normalized intensity of LAMP1-msGFP (cyan) and JF549i (yellow). The labels i and ii are labeled on the line profile as well and correspond to the same labels in h). Cartoon schematic of analyzed signal is above graph.

The IC_50_ for DAPT inhibition of γ-secretase mediated cleavage of SYT7 was ~71 nM (Fig. 5b-c), which is similar to the IC_50_ of another γ-secretase target, amyloid precursor protein (APP) (**Dovey et al., 2001**). To address the location of SYT7 processing by γ-secretase, we transduced neurons with HaloTag-SYT7α and added an impermeant, nonfluorogenic, JaneliaFluor dye (JF549i) to the media at a low concentration (1 nM) for 2 days. HaloTag-SYT7α transiting through the plasma membrane before cleavage (processing by γ-secretase), will become labelled with extracellular JF549i, allowing us to follow the intracellular fluorescent adduct. However, if SYT7 is processed at or before it reaches the plasma membrane, fluorescence will not be observed (Fig. 5d). Indeed, full length SYT7 is present on the plasma membrane and is apparently cleaved in synaptic endosomal structures because we observe small JF549i puncta (yellow) throughout the soma that partially colocalizes with LAMP1 (cyan) positive structures (Fig. 5e). Interestingly, these punctae localize to the lumen (i) of LAMP1 positive structures, or to a portion of the endo-lysosomal membrane (ii), or do not colocalize with LAMP1 at all (iii) (Fig. 5f). As a control, JF549i did not label untransduced neurons, so all labelling was specific for tagged SYT7α (Fig. S5e).

### SYT7 is mislocalized and destabilized when amino-terminal cleavage is blocked

It remained unclear whether γ-secretase processing supports the axonal localization or function of SYT7, or whether γ-secretase processing is part of the normal degradation pathway for this protein. Interestingly, SYT7 is palmitoylated near its TMD and this post-translational modification has been shown be important for lysosomal trafficking of SYT7 in fibroblasts (**Flannery et al., 2010**). Here we examined SYT7 localization in control neurons and neurons treated with: DAPT, 2-bromopalmitate (2BP), and DAPT + 2BP treatments; 2-BP is a palmitoylation inhibitor (**Webb, Hermida-Matsumoto, & Resh, 2000**). Proteins with palmitoylation sites are dynamically de-palmitoylated and re-palmitoylated; adding a palmitoylation inhibitor biases the protein to a de-palmitoylated state. For these experiments, we included SYT1 as a control. SYT1 is responsible for fast, synchronous SV fusion, and like SYT7, it is palmitoylated in or near its lone TMD (**Chapman et al., 1996; Chapman, 2008**). We also included LAMP1-msGFP as another general, membrane anchored protein control; this construct also allowed us to further examine the colocalization of SYT7α and LAMP1+ structures.

Neurons were untreated (control), treated with DAPT for 10-12 days, with 2BP for 3 hrs, or both, and then fixed and stained for SYT1, SYT7, and the axon initial segment (AIS). None of these treatments affected the localization of SYT1 (Fig. 6a), whereas DAPT treatment resulted in the mislocalization of the majority of SYT7α-HaloTag to small punctae that appear similar to the structures shown in Fig. 5e, only faint axonal staining was apparent during DAPT treatment. Surprisingly, short treatment with 2BP led to the complete absence of SYT7α-HaloTag, as did the combination of DAPT and 2BP (Fig. 6b). Accordingly, substitution of the palmitoylated cysteine residues results in an unstable protein that is only marginally stabilized upon DAPT treatment (Fig. S6b). These results reveal that palmitoylation plays an essential role in determining SYT7 stability. The punctate SYT7α-HaloTag positive structures observed during DAPT treatment appeared at the detriment of normal axonal and lysosomal localization (Fig. 6c-d); under these conditions, SYT7α-HaloTag mislocalizes to the earlier secretory pathway at the expense of the later secretory pathway (Fig. 6d). However, treatment of WT neurons with DAPT does not phenocopy the SYT7KO phenotype (Fig. S6c-e). This lack of an effect might result from low levels of axonal membrane targeted SYT7, even with DAPT treatment. SYT7 is a long-lived presynaptic protein, so we next investigated whether γ-secretase processing influences its half-life (**Dorrbaum, Kochen, Langer, & Schuman, 2018**). We found that SYT7 is indeed a long-lived protein and that γ-secretase inhibition enhances its turnover (Fig. 6e-f and S6a). This was somewhat unexpected because γ-secretase processing is conventionally thought to accelerate the turnover of its substrates (**Kopan & Ilagan, 2004**). In summary, SYT7 is cleaved at the amino terminus by γ-secretase which is critical to maintain axon membrane localization and protein half-life.

**Figure 6:**
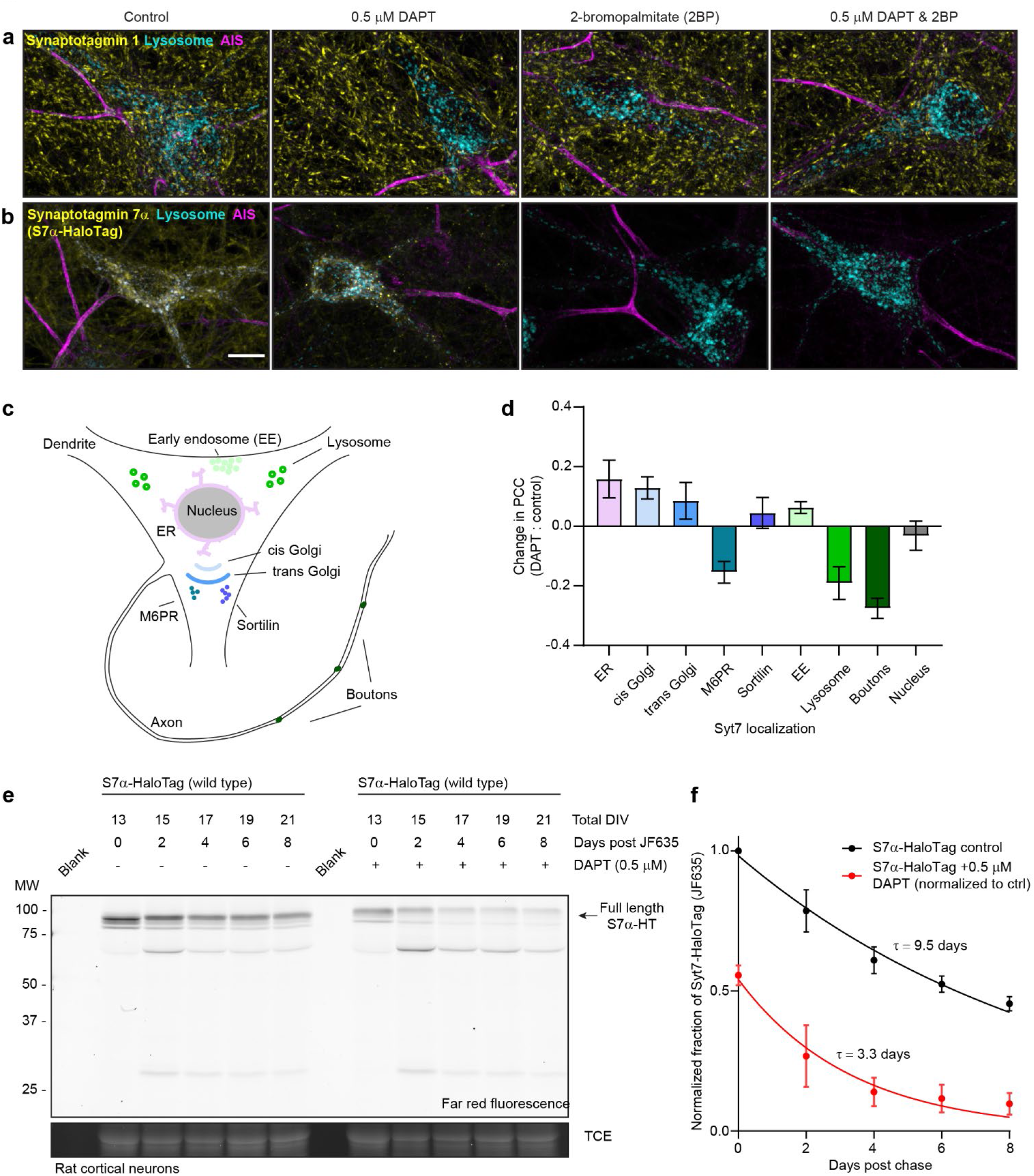
SYT7 is mislocalized and destabilized when amino-terminal cleavage is blocked. **a**) Representative super-resolution maximum z-projections of rat hippocampal neurons transduced with LAMP1-msGFP (cyan), fixed for ICC, and stained for SYT1 (yellow) and the AIS (magenta). Four separate conditions were imaged, control neurons, neurons grown for 10-12 days in 0.5 μM DAPT, neurons exposed to 2BP for 3 hours before imaging, and neurons exposed to the combination of DAPT and 2BP treatment. **b**) Same as in a), but instead of anti-SYT1 staining, neurons were transduced with SYT7α-HaloTag and reacted with JF549 during overnight primary antibody incubation to monitor SYT7α localization. Scale bar 10 microns. **c**) Illustration of model neuron and compartments assayed for SYT7α-HaloTag colocalization. **d**) Bar graph showing changes in colocalization of SYT7α-HaloTag/JF549 and labelled organelles (M6PR and Sortilin label post-Golgi vesicles). Quantified by taking the difference of Pearson’s R (PCC) between DAPT treated and control neurons in each condition. Values are means +/− error propagated SEM from at least 3 separate experiments for each condition. **e**) Representative in-gel fluorescence of protein extracted from rat cortical neurons transduced with SYT7α-HaloTag and pulse-chased with JF635 at 13 DIV under control conditions and when grown in 0.5 μM DAPT. Cultures were labeled with JF635 at 13 DIV and then robustly washed with conditioned media. The disappearance of labeled SYT7α-HaloTag/JF635 from the gel, can be used to calculate protein half-life. Control SYT7α-HaloTag/JF635 runs between 75 and 100 kDa, while DAPT treated SYT7α-HaloTag/JF635 runs slightly higher because cleavage of the amino terminus is blocked. TCE is used as a load control. **f**) Normalized intensity of SYT7α-HaloTag/JF635 plotted as fraction of total control SYT7α-HaloTag/JF635 against days post wash. Values are means +/− SEM from 3 independent experiments. Single exponential decay functions were fitted to control (black) and DAPT (red) conditions. The tau for control SYT7α-HaloTag/JF635 is 9.5 days, while the tau for DAPT treated SYT7α-HaloTag/JF635 is 3.3 days.

### Dissociating discrete SYT7 functions via protein retargeting

We demonstrated above that SYT7 influences PPF, asynchronous release, and SV replenishment, and its cellular location and stability is regulated by γ-secretase processing and palmitoylation. We therefore asked how the location of SYT7 influences the SV cycle. To our surprise, the distinct functions of SYT7 in the SV cycle could be dissociated by retargeting the protein to different destinations. For these experiments, we restricted SYT7α to the plasma membrane (PM), endo-lysosomal LAMP1+ membranes, or SVs, by replacing the luminal amino terminus and TMD from SYT7α with different targeting motifs. To target SYT7α to the PM we added a preprolactin signal sequence followed by a CD4 TMD and a Golgi export sequence (Fig. 7a). For endo-lysosomal membrane targeting, fusing the cytosolic portion of SYT7α to the carboxy terminus of LAMP1 was sufficient (Fig. 7b). Similarly, for targeting to SVs, we fused the cytosolic domain of SYT7α to SYP (Fig. 7c and Fig. S7a). Note, in Fig. 7a-c, retargeted SYT7α constructs were sparsely transduced to clearly demonstrate cellular localization. When expressed in HEK293T cells, these constructs also localized to the PM, endo-lysosomal compartment, and small vesicles, respectively (Fig. S7b). Interestingly, SYT7α that was restricted to the PM by replacing the WT TMD with a CD4 TMD, and adding a viral Golgi export sequence, demonstrated a polarized distribution along axons. Therefore, γ-secretase processing is an essential prerequisite for enrichment of WT SYT7α to the axon PM, but that there is another axonal targeting motif in the protein (Fig. 7a inset and Fig. S7c).

**Figure 7:**
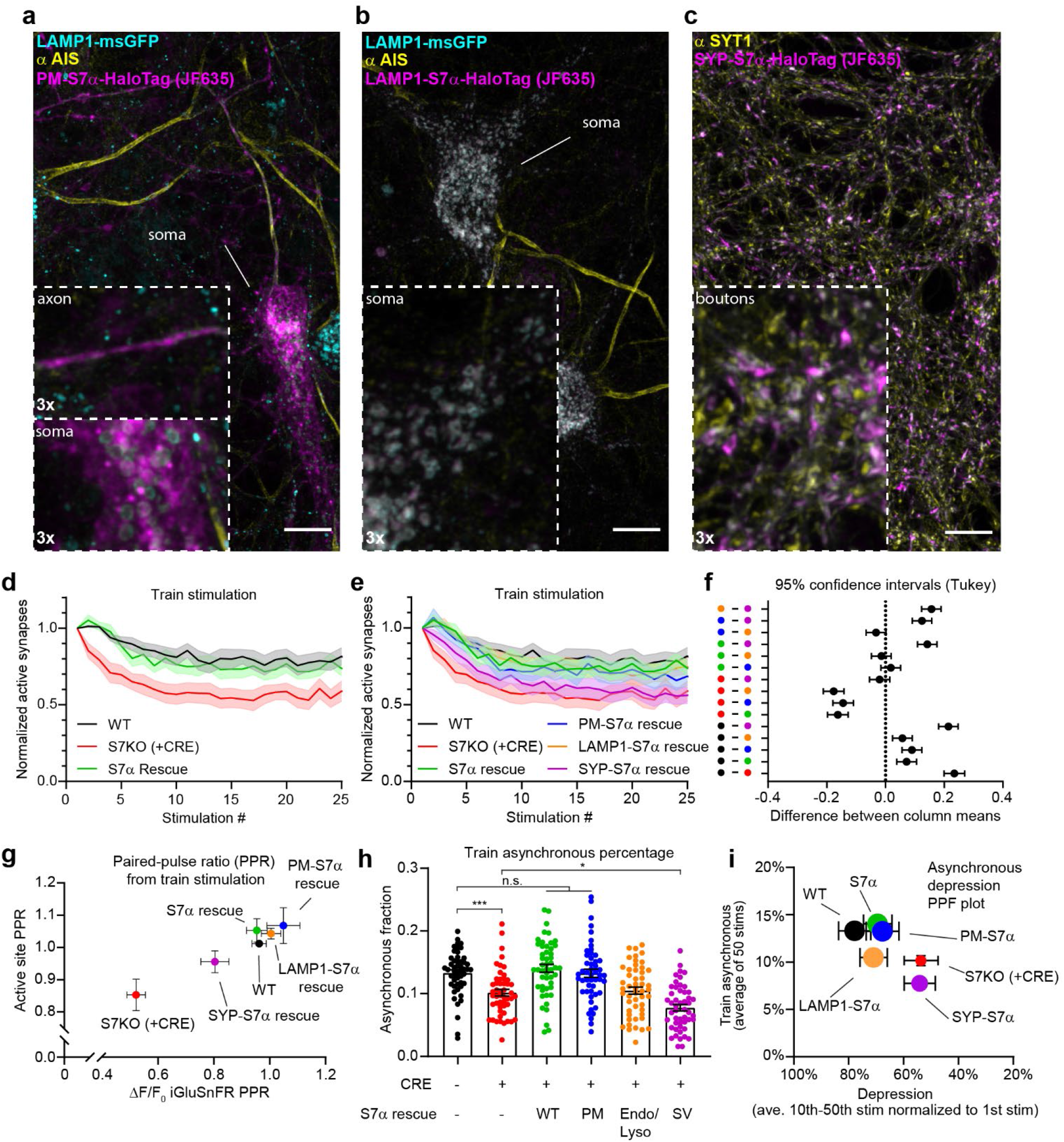
Dissociating discrete SYT7 functions via protein retargeting. **a**) Representative super-resolution maximum z-projection of rat hippocampal neurons transduced with LAMP1-msGFP (cyan) and PM-SYT7α-HaloTag (magenta) (a plasma membrane targeted SYT7α), fixed and stained with JF635 and anti-pan-neurofascin (yellow) antibodies. The 3x zoom inset shows axonal localization (top) of PM-SYT7α, with little to no detectable localization on soma lysosomal structures (bottom). **b**) Representative image taken with similar conditions as in a), but instead of transducing with PM-SYT7α-HaloTag, LAMP1-SYT7α-HaloTag was transduced. The 3x zoom inset shows exclusively lysosomal localization of this SYT7α construct. **c**) Representative image taken with similar conditions as in a) and b) but transduced with SYP-SYT7α-HaloTag. The 3x zoom inset shows SYP-SYT7α-HaloTag localization to the presynapse. a)-c) Rescue SYT7α-HaloTag constructs for ICC were sparsely transduced to better examine localization. Scale bars = 10 microns. **d**) Depression plot, fraction of active synapses (synapses releasing peak glutamate above baseline, >4 SD above noise) as a function of stimulation number during HFS. Values are means (solid line) +/− SEM (shaded error), WT (black, n = 15), SYT7KO (red, n = 13), and SYT7α rescue (green, n = 15) from 3 independent experiments, SYT7KO vs SYT7α rescue is p < 0.0001 by two-way ANOVA comparing genotype. **e**) Depression plot from d), but with rescue SYT7α constructs included. Values are means (solid line) +/− SEM (shaded error), PM-SYT7α rescue (blue, n = 15), LAMP1-SYT7α rescue (orange, n = 15), and SYP-SYT7α rescue (purple, n = 15) from 3 independent experiments. **f**) Multiple comparison confidence interval (95% CI) plot from dataset in e). Plot generated from two-way ANOVA comparing the predicted mean difference between genotypes of normalized active synapses. Comparisons with errors encompassing zero are not statistically different. Total summary statistics are included in supplementary statistics table 6. **g**) An X-Y plot of paired-pulse ratio generated at 20 Hz (from first two pulses of HFS). Values are means +/− SEM where X values are the ratio of the change in glutamate release (ΔF/F_0_ iGluSnFR peaks) and Y values are the fraction of ROIs releasing glutamate (active sites) from WT (black), SYT7KO (red), and SYT7α rescue (green), PM-SYT7α rescue (blue), LAMP1-SYT7α rescue (orange), and SYP-SYT7α rescue (purple). **h**) Train asynchronous release (peak release recorded between 10 ms and 50 ms post stimulus) of WT and SYT7KO vs the labeled rescue constructs. Values are means +/− SEM and are the average asynchronous values from each stimulus during a 50 A.P. (20 Hz) HFS, so n = 50 for each group. All comparisons and summary statistics are provided in supplementary statistics table 7, only some are labeled on the graph for presentations sake, p-values are as follows *** = 0.001, * = 0.0147, by one-way ANOVA with Holm-Sidak’s multiple comparisons test. **i**) Summary X-Y plot illustrating different magnitudes of rescue for three of the proposed functions of SYT7. Values are means +/− SEM where X values represent depression percentage (release from 10^th^-50^th^ stimulation normalized to 1^st^), Y values are the average asynchronous percentage of each genotype during the 50 A.P. of a HFS, and the size of each mark is the relative size of each’s PPR, normalized on a scale from the largest - 10 (most PPF) to smallest - 1 (least PPF).

To examine the function of these rescue constructs, we chose to use HFS so that we could measure: 1) 20 Hz PPF, 2) train related asynchronous release, and 3) synaptic depression and SV replenishment. For these experiments we used a new floxed SYT7 mouse line (MRC Harwell Institute #Syt7-TM1C-EM4-B6N). This inducible KO avoids any developmental confounding factors due to chronic inactivation of SYT7, while at the same time serves to reduce animal waste. These experiments also validate our SYT7KO phenotypes in a separate genetic line and establish this new SYT7 floxed line for future use (Fig. S7d-e). The expression of all rescue constructs was confirmed via immunoblot analysis (Fig. S7f). First, we found that expression of untagged SYT7α rescues synaptic depression (Fig. 7d). We further observed that the PM and endo-lysosomal targeted constructs both also rescue synaptic depression, while SV-targeted SYT7α, does not (Fig. 7e-d). This is rather remarkable because SV targeted SYT7α is present at the site of exocytosis; the observation that this construct does not rescue the KO phenotype emphasizes the importance of precise SYT7α localization. Confidence intervals for the difference between total active synapses throughout the stimulus train are shown in Fig. 7f, which provides a compact means to visualize all pair-wise comparisons. By quantifying the first two stimuli from the HFS experiments, we calculated the 20 Hz PPF ratio. Using the same methods as in Fig. 1e-f, we plotted the two components of facilitation with active synaptic sites on the y-axis and peak iGluSnFR changes in fluorescence on the x-axis. Here we see that the WT PPF ratio is positive and clusters with full length SYT7α, PM-, and LAMP1-SYT7α rescue constructs; in contrast SV-targeted SYT7α failed to rescue PPF. Examining asynchronous release over a train, both SYT7α and PM-SYT7α rescue asynchronous release in SYT7KO neurons, but the LAMP1-SYT7α construct does not, even though it rescues other release modes. Strikingly, when targeted to SVs, SYT7α unexpectedly promoted synchronous release instead of asynchronous release (Fig. 7h). We also plotted all conditions tested as the synchronous fraction against stimulation number (Fig. S7g), where best fit lines are shown for clarity. This plot illustrates that as the number of successive stimuli increase, the fraction of asynchronous release increases as well.

To summarize the three phenotypes described here, with respect to the rescue constructs, we plotted the average asynchronous fraction of release on the y-axis, depression is plotted on the x-axis, and the PPF ratio is encoded in the size of each point in the graph (largest size is the highest PPF ratio, smaller size represents lower, or negative PPF ratio) (Fig. 7i). As we hypothesized earlier, the location of SYT7α is paramount to its function. Here we untangle the various functions of SYT7α through restricting and retargeting techniques. Only the plasma membrane associated SYT7α rescued all investigated phenotypes. Interestingly, the SV retargeted SYT7α not only failed to rescue asynchronous release but did the opposite, promoting synchronous release. While the underlying mechanism is unclear, this construct provides a novel tool to tune synchronous release at central synapses.

## Discussion

SYT7 is highly expressed in the brain but a consensus regarding its precise function in neurons remains the subject of considerable debate. Multiple first reports, examining synaptic function using electrophysiology and constitutive KO mouse lines, came to the same conclusion: SYT7 played no role in synaptic transmission or the SV cycle (**Maximov et al., 2008**). Later, upon the application of extended stimulus trains, deficiencies in SYT7KO neurons were reported; these included reductions in asynchronous release (**Wen et al., 2010**) or enhanced synaptic depression (**Liu, Bai, Hui, et al., 2014**). Recent studies indicate that the role of SYT7 in the SV cycle may depend on the type of synapse that is under study (**Jackman et al., 2016; Turecek & Regehr, 2018**). Here, we took a different approach. We chose to use dissociated mouse hippocampal neurons because they are a ubiquitous model system in the field, and this preparation allows for tractable investigation of the underlying molecular and cellular mechanisms. Additionally, we applied state-of-the-art molecular tools to probe SYT7 function including optical reporting of presynaptic glutamate release (iGluSnFR), millisecond and nanometer resolution of the SV cycle (zap-and-freeze EM), and enhanced protein detection using various HaloTag ligands (JaneliaFluor probes). We show that the three debated SV cycle phenotypes reported at SYT7KO synapses all occur, to varying degrees, at hippocampal synapses from dissociated cultures. Based on our data, we argue that wherever SYT7 is present, it supports PPF, asynchronous release, and SV replenishment.

Using iGluSnFR we detected a small change in asynchronous release from single stimuli between WT and SYT7KO synapses (Fig. 1), but otherwise, there was no apparent change in the amplitude of release from a single stimulus. During PPF measurements, we observed that SYT7KO neurons failed to facilitate. Optical detection of release allowed us to further explore the nature of facilitation. In WT synapses, PPF is due to enhanced glutamate release from already active synapses, and not from an additional activation of previously silent synapses (i.e. recruitment). In SYT7KO neurons, glutamate release is reduced (turning to depression), and the number of active synapses decreases. Additionally, measuring release during HFS via conventional whole cell patch clamp produces a train of responses that fail to decay to baseline. This charge transfer component is termed tonic transmission and was thought to arise due to an accumulation of glutamate during HFS, from the spillover of glutamate, or from asynchronously released glutamate. Using iGluSnFR to monitor release during HFS, we measure glutamate release from individual synapses, and we argue that ‘tonic’ transmission results from an increasing fraction of asynchronously released SVs as the stimulus train progresses. We suggest that this is an activity-dependent form of more slowly released SVs, and that this mode of asynchronous release is decreased at SYT7KO synapses (Fig. 2). Importantly, our reasoning follows from comparing averaged iGluSnFR traces to individual iGluSnFR ROIs; in individual traces, during steady-state release, iGuSnFR signals from individual ROIs decay to baseline, whereas averaged iGluSnFR signals do not, strongly supporting asynchronous release as a driver of increased baseline fluorescence. However, because we are employing the low affinity iGluSnFR, there may be ‘residual’ glutamate that electrophysiological measurements detect, but iGluSnFR does not.

Having demonstrated that release is universally reduced in SYT7KO neurons after an initial stimulus (asynchronous, PPF, depression), we sought to investigate how this is manifested in the SV cycle using ‘zap-and-freeze’ EM. This approach revealed that docked vesicles are more severely depleted by both single stimuli and HFS in SYT7KO (Fig. 3). Hence, SYT7 serves to promote vesicle docking, or to preventing undocking after a single stimulus and during a stimulus train (Fig. 3). Interestingly, we were also able to document decreases in the total number of SVs during HFS that, in WT neurons, recovered within 5 s, but in the SYT7KO, failed to completely recover over the same time frame. This decreased SV number phenotype seen after HFS may partly explain the replenishment defect observed here (Fig. 2) and elsewhere (**Liu, Bai, Hui, et al., 2014; Chen et al., 2017**). By combining optical SV exocytosis methods with ‘zap-and-freeze’, we can pin-point where the defect arises in SYT7KO synapses during the SV cycle. This leads to a paradox however, because the docking defect and the failure to recover SV number presumably occur in two different areas of the presynaptic zone. We therefore examined the trafficking and localization of SYT7 which led us to the observation that SYT7 is processed by the γ-secretase complex (Fig. 5).

Membrane protein processing by the γ-secretase complex, while originally postulated to be an intramembrane proteasome, is now thought to regulate the location of membrane proteins (**Kopan & Ilagan, 2004**). The cleavage of SYT7 by γ-secretase, along with this protein’s high sensitivity to palmitoylation inhibitors, may afford SYT7 with the ability to quickly transfer between membranes in the presynapse during sustained stimulation, irrespective of the sorting of SV proteins. Support for this idea stems from: 1) the speed with which SYT7 is depalmitoylated in the presence of 2-BP, a palmitoylation inhibitor, which suggests active and robust palmitoylation/depalmitoylation cycling, 2) γ-secretase cleavage stabilizes SYT7, and 3) γ-secretase cleavage promotes axonal localization of WT SYT7 (Fig. 6). Support against this idea comes from our WT inhibitor and rescue experiments. If γ-secretase cleavage is absolutely needed for SYT7 function, we should phenocopy the SYT7KO by applying inhibitors to WT neurons (Fig. S6), but this was not observed. However, other experiments show that SYT7 must transit through the plasma membrane and that during inhibitor (DAPT) treatment, a detectable amount of SYT7 was still present on axons. So, it is plausible that there is enough SYT7 at the PM to sustain synaptic function in the γ-secretase inhibitor experiments. Interestingly, in synapses lacking components of the γ-secretase complex, namely presenilin, have strikingly similar SV cycle phenotypes to SYT7KO synapses, specifically enhanced depression and reduced PPF (**Zhang et al., 2009**). The PM restricted SYT7 rescued all functions of WT SYT7; since this construct includes a viral Golgi export sequence to enhance PM targeting, it remains possible that this sequence supplants the role of γ-secretase cleavage and palmitoylation.

While a portion of untagged SYT7 localizes to the endo-lysosomal compartment, and the retargeted construct (LAMP1-SYT7α) rescued some of the functions of SYT7, we could not define an absolute role for SYT7, on lysosomes, in the SV cycle. Moreover, no alterations in lipids or cholesterol sensitive SV protein interactions were observed. We examined this issue due to reports concerning the role of SYT7 in maintaining lysosome-peroxisome contacts and in cholesterol trafficking in fibroblasts (**Chu et al., 2015**). Interestingly, PM and endo-lysosomal targeted SYT7α were both able to rescue the synaptic depression and paired-pulse facilitation phenotypes that we observed; however, the endo-lysosomal targeted SYT7α did not rescue asynchronous release (Fig. 7). This is an important observation because paired-pulse facilitation and asynchronous release could have shared a common mechanism; if so, they would not be separable. By retargeting SYT7α, we found that these functions could be disassociated from one another, so these processes are likely to be mechanistically distinct. It should be noted that small amounts of LAMP1 can be detected on the PM, but we did not detect restricted endo-lysosomal SYT7 on the PM in our images, so our approach appears to be valid. More striking was the observation that when targeted to SVs, SYT7α did not rescue any SYT7KO synaptic phenotype and instead suppressed asynchronous release, opposite to its normal function. When Double C2-like domain-containing protein (DOC2), another protein that plays a role in asynchronous release, was tethered to SVs, it enhanced this slow phase of transmission (**Yao, Gaffaney, Kwon, & Chapman, 2011**). Future studies will address why syt7 does not behave in the same manner, but the retargeted protein already provides a useful tool to modulate the extent of asynchronous release to potentially alter, for example, reverberatory activity (**Lau & Bi, 2005**).

In summary, we have shown that SYT7 participates in asynchronous release, short-term synaptic plasticity, and SV replenishment in individual hippocampal synapses. Notably, we have provided for the first time, SV morphological correlates that help to explain the observed defects in SYT7KO synapses: SVs in SYT7KO synapses undock to a greater extent than WT controls and fail to efficiently regenerate SVs following high frequency stimulation. In the quest of the location-function correlation of SYT7, we have identified SYT7 as a substrate for the γ-secretase complex and dissociated the roles of SYT7 in asynchronous release and paired-pulse facilitation. These findings lead to a new integrated view of the role of SYT7 in the SV cycle (summary figure). Future work will involve careful ‘zap-and-freeze’ experiments to follow endosomal intermediates produced during HFS. New advances in correlative light and electron microscopy (CLEM) or fluorescence electron microscopy (fEM) may make the identification of the endosomal intermediates that are influenced by SYT7 possible. Moreover, the role of SYT7 and DOC2 isoforms in promoting asynchronous release during train stimulation needs further clarification. Clearly, both proteins influence asynchronous release but to what exact extent and how they may interact remains unresolved. In the experiments reported here, SYT7 promoted late train asynchronous release so it is tempting to speculate that both proteins promote asynchronous release but during different phases of the stimulation epoch; perhaps early asynchronous release is DOC2 dependent while late asynchronous release is SYT7 dependent. One model that we will explore is whether SYT7 functions mainly as a promoter of docking and priming while DOC2 mediates the actual triggering of asynchronously released vesicles.

## Acknowledgements

This study was supported by grants from the NIH (MH061876 and NS097362 to E.R.C.). J.D.V. was supported by a postdoctoral fellowship from the National Institutes of Health F32 NS098604. E.R.C. is an Investigator of the Howard Hughes Medical Institute. SW was supported by start-up funds from the Johns Hopkins University School of Medicine, Johns Hopkins Discovery funds, Johns Hopkins Catalyst Award, the National Science Foundation (1727260), and the National Institutes of Health (1DP2 NS111133-01 and 1R01 NS105810-01A1). S.W. is an Alfred P. Sloan fellow, a McKnight Foundation Scholar and a Klingenstein and Simons Foundation scholar. G.F.K. was supported by a grant from the National Institutes of Health to the Biochemistry, Cellular and Molecular Biology program of the Johns Hopkins University School of Medicine (T32 GM007445) and is a National Science Foundation Graduate Research Fellow (2016217537). The EM ICE high-pressure freezer was purchased partly with funds from an equipment grant from the National Institutes of Health (S10RR026445) awarded to S. C. Kuo.

## Author Contributions

J.D.V. and E.R.C. conceived of the project. J.D.V., G.F.K., K.C.C., S.W. and E.R.C. and designed the experiments. J.D.V., G.F.K., E.C. and K.C.C. performed the experiments. J.D.V., G.F.K. and S.W. analyzed the data. S.W. and E.R.C. supervised the projects. J.D.V. and E.R.C. wrote the manuscript.

## Methods

### Ethics Statement

Animal care and use in this study were conducted under guidelines set by the NIH Guide for the Care and Use of Laboratory Animals handbook. The protocols were reviewed and approved by the Animal Care and Use Committee (ACUC) at the University of Wisconsin, Madison (Laboratory Animal Welfare Public Health Service Assurance Number: A3368-01).

### Cell Culture

Sprague Dawley rat hippocampal and cortical neurons were isolated at E18 (Envigo). Mouse hippocampal neurons from the *syt7* floxed mouse strain Syt7-TM1C-EM4-B6N (**Codner et al., 2018**) and *syt7^tm1Nan^* (**Chakrabarti et al., 2003**) were isolated at P0 and prepared using a procedure previously described in (**Vevea & Chapman, 2020**). Briefly, hippocampal neurons were dissected, trypsinized (Corning; 25-053-CI), triturated, and plated on glass coverslips (Warner instruments; 64-0734 (CS-18R17)) coated with poly-D-lysine (Thermofisher; ICN10269491) and EHS laminin (Thermofisher; 23017015). Neurons were grown in Neurobasal-A (Thermofisher; 10888-022) medium supplemented with B-27 (2% Thermofisher; 17504001), Glutamax (2 mM Gibco; 35050061), and pen/strep before experiments. For high-pressure freezing and electron microscopy (EM), cell cultures were prepared on 6-mm sapphire disks (Technotrade), mostly as previously described (**Kusick et al., 2020**). For two of the three experiments/cultures, genotyping was performed after hippocampal dissection, using cortices, with hippocampi left in NB-A before switching to papain after, while in the third genotyping was performed using tail clips. Before use, sapphire disks were carbon-coated with a “4” to indicate the side that cells are cultured on. Health of the cells, as indicated by de-adhered processes, floating dead cells, and excessive clumping of cell bodies, was assessed regularly, as well as immediately before experiments. All EM related experiments were performed between 13 and 17 DIV. All other experiments were performed between 14 to 23 DIV. For virus preparation, HEK293T cells (ATCC) were cultured following ATCC guidelines and were tested for mycoplasma contamination using the Universal Mycoplasma Detection Kit (ATCC; 30-1012K); HEK293T cells were validated using Short Tandem Repeat profiling by ATCC (ATCC; 135-XV) within the previous year.

### Lentivirus production and use

Lentivirus production was performed as described previously (**Vevea & Chapman, 2020**). Lentiviral constructs were all subcloned into the FUGW transfer plasmid (FUGW was a gift from David Baltimore (Addgene plasmid # 14883; http://n2t.net/addgene:14883; RRID:Addgene_14883)) (**Lois, Hong, Pease, Brown, & Baltimore, 2002**). We previously replaced the ubiquitin promoter with the CAMKII promoter or human synapsin I promoter (**Kugler, Kilic, & Bahr, 2003; Vevea & Chapman, 2020**). Lentivirus that expresses CRE was added to neuronal cultures at 1 DIV; viruses that express iGluSnFR or pHluorin constructs were added at 1 and 5 DIV respectively. SYT7α rescue and other constructs that were used to mark organelles, were added at 5-6 DIV.

### Plasmid construction

Two types of plasmids were used in this study. One was our previously modified lentivirus backbone of choice derived from FUGW (**Vevea & Chapman, 2020**), and the other was based off pEF-GFP, excising the GFP and substituting our own various inserts (pEF-GFP was a gift from Connie Cepko (Addgene plasmid # 11154; http://n2t.net/addgene:11154; RRID:Addgene_11154)) (**Matsuda & Cepko, 2004**). The low affinity glutamate sensor iGluSnFR S72A was PCR amplified from the pAAV.hSynapsin.SF-iGluSnFR.S72A which was a gift from Loren Looger (Addgene plasmid # 106176; https://www.addgene.org/106176/; RRID:Addgene_106176)) (**Marvin et al., 2018**), and subcloned into our lentivirus transfer plasmid (CamKII promoter) along with the addition of membrane trafficking motifs to promote plasma membrane localization as done previously for the original iGluSnFR variant (**Vevea & Chapman, 2020**). The vGlut1-pHluorin construct used here was subcloned into our modified FUGW transfer vector (hSynapsin promoter) from the original vGlut1-pHluorin construct (**Voglmaier et al., 2006**). The synaptotagmin 7a rescue constructs were assembled using PCR SOE and subcloned into our modified FUGW transfer plasmid. To retarget SYT7a, we fused the cytosolic domain (including juxtamembrane linker), to various protein domains. For plasma membrane (PM) targeted SYT7a we used a preprolactin signal sequence (PP) fused to a CD4 transmembrane domain (TMD) along with an adjacent Golgi export sequence (**Parmar, Barry, Kai, & Duncan, 2014**) amended to the amino terminus of SYT7a. For endo-lysosomal targeting we added the cytosolic domain of SYT7a to the carboxy terminus of the LAMP1 protein, and for synaptic vesicle targeting, the cytosolic domain of SYT7α was fused to the carboxy terminus of synaptophysin. For HaloTag fusions, the HaloTag cassette was amplified from the pHTC HaloTag® CMV-neo Vector (Promega; G7711) and amended to either the amino or carboxy terminus of SYT7a constructs. If added to the amino terminus, a preprolactin leader sequence was included to ensure protein translocation across the ER membrane. The fluorescent proteins msGFP (Vevea et al 2020) or mRuby3 (**Bajar et al., 2016**) were appended to the carboxy terminus of LAMP1 for use as a lysosomal marker, synaptophysin as a synaptic vesicle marker, the plasma membrane motifs mentioned above for plasma membrane targeting, or alone for a cytosolic marker. For CRE expression, we used pLenti-hSynapsin-CRE-WPRE (pLenti-hSynapsin-CRE-WPRE which was a gift from Fan Wang (Addgene plasmid # 86641; http://n2t.net/addgene:86641; RRID:Addgene_86641)) (**Sakurai et al., 2016**). All original plasmids used in this study are deposited in Addgene, filed under this manuscript.

### SYT7 constructs

pF(UG) hSyn Syt7a
pF(UG) hSyn PP-HaloTag-Syt7a
pF(UG) hSyn Syt7a-HaloTag
pF(UG) hSyn Syt7a-HaloTag TMD Cys-Ala
pF(UG) hSyn Syt7a-HaloTag P2A PM-msGFP
pF(UG) hSyn SYP-∆TMD Syt7a
pF(UG) hSyn SYP-∆TMD Syt7a-HaloTag
pF(UG) hSyn PM-Syt7a∆TMD
pF(UG) hSyn PM-Syt7a∆TMD-HaloTag
pF(UG) hSyn Lamp1-Syt7a∆TMD
pF(UG) hSyn Lamp1-Syt7a∆TMD-HaloTag

### Biosensors

pF(UG) CamKII sf iGluSnFR S72A
pF(UG) hSyn vGlut1 pHluorin

### Organelle markers

pF(UG) hSyn SYP-mRuby3
pF(UG) hSyn Lamp1-msGFP (JV012)
pEF-GFP
pEF mRuby3

### Immunoblot protocol

Immunoblots were performed as described previously, (**Vevea & Chapman, 2020**). Primary antibodies were: anti-SYT1 (1:1000, 48) (lab stock; mAB 48; RRID:AB_2199314), anti-SYP (1:1000) (SySy; 101 004; RRID:AB_1210382), anti-SYT7 (1:1000) (SySy; 105 173; RRID:AB_887838), anti-HaloTag® (1:1000) (Promega; G9211; RRID:AB_2688011). Secondary antibodies were: goat anti-mouse IgG2b-HRP (Biorad, M32407; RRID:AB_2536647), goat anti-mouse IgG-HRP (Biorad, 1706516; RRID:AB_11125547), goat anti-rabbit IgG-HRP (Biorad, 1706515, RRID:AB_11125142), goat anti-guinea pig IgG-HRP (Abcam, ab6908, RRID:AB_955425).

### High-pressure freezing and freeze-substitution

Cells cultured on sapphire disks were frozen using an EM ICE high-pressure freezer (Leica Microsystems), exactly as previously described (**Kusick et al., 2020**). The freezing apparatus was assembled on a table heated to 37 °C in a climate control box, with all solutions pre-warmed (37 ºC). Sapphire disks with neurons were carefully transferred from culture medium to a small culture dish containing physiological saline solution (140 mM NaCl, 2.4 mM KCl, 10 mM HEPES, 10 mM glucose; pH adjusted to 7.3 with NaOH, 300 mOsm). NBQX (3 μM; Tocris) and bicuculline (30 μM; Tocris) were added to the physiological saline solution to block recurrent synaptic activity. CaCl_2_ and MgCl_2_ concentrations were 1.2 mM and 3.8 mM, respectively. After freezing, samples were transferred under liquid nitrogen to an EM AFS2 freeze substitution system at −90 °C (Leica Microsystems). Using pre-cooled tweezers, samples were quickly transferred to anhydrous acetone at −90 °C. After disassembling the freezing apparatus, sapphire disks with cells were quickly moved to cryovials containing freeze substitution solutions. For the first two experiments, freeze substitution was performed exactly as previously described^3^: solutions were 1% glutaraldehyde, 1% osmium tetroxide, and 1% water in anhydrous acetone, which had been stored under liquid nitrogen then moved to the AFS2 immediately before use. The freeze substitution program was as follows: −90 °C for 6-10 hr (adjusted so substitution would finish in the morning), 5 °C h^−1^ to −20 °C, 12 h at −20 °C, and 10 °C h^−1^ to 20 °C. For the third experiment, we used a different freeze substitution protocol that yields more consistent, high-contrast morphology: samples were first left in 0.1% tannic acid and 1% glutaraldehyde at −90 °C for ~36 hours, then washed 5x, once every 30 min, with acetone, and transferred to 2% osmium tetroxide, then run on the following program: 11 hr at −90 °C, 5 °C h^−1^ to −20 °C, −20 °C for 12 hr, 10 °C h^−1^ to −4 °C, then removed from the freeze substitution chamber and warmed at room temperature for ~15 min before washing.

### Embedding, sectioning, and transmission electron microscopy

Embedding and sectioning were performed exactly as previously described (**Kusick et al., 2020**). For ultramicrotomy, 40-nm sections were cut. Sections on single-slot grids coated with 0.7% pioloform were stained with 2.5% uranyl acetate then imaged at 80 kV on the 93,000x setting on a Phillips CM 120 transmission electron microscope equipped with an AMT XR80 camera run on AMT Capture v6 for the first experiment, and on a for the other two experiments, samples were imaged on a Hitachi 7600 TEM equipped with an AMT XR50 run on AMT Capture v6. Samples were blinded before imaging. To further limit bias, synapses were found by bidirectional raster scanning along the section at 93,000x or 100,000x, which makes it difficult to “pick” certain synapses, as a synapse usually takes up most of this field of view. Synapses were identified by a vesicle-filled presynaptic bouton and a postsynaptic density. Postsynaptic densities are often subtle in our samples, but synaptic clefts were also identifiable by 1) their characteristic width, 2) the apposed membranes following each other closely, and 3) vesicles near the presynaptic active zone. 125-150 micrographs per sample of anything that appeared to be a synapse were taken without close examination.

### Electron microscopy image analysis

Electron microscopy image analysis was performed as previously described (**Kusick et al., 2020**). All the images from a single experiment were randomized for analysis as a single pool. Only after this randomization were any images excluded from analysis, either because they appeared to not contain a *bona fide* synapse or the morphology was too poor for reliable annotation. The plasma membrane, active zone, docked SVs, and all SVs in the bouton were annotated in ImageJ using SynapsEM plugins (**Watanabe, Davis, Kusick, Iwasa, & Jorgensen, 2020**) (https://github.com/shigekiwatanabe/SynapsEM). The active zone was identified as the region of the presynaptic plasma membrane with the features described above for identifying a synapse. Docked vesicles were identified by their membrane appearing to be in contact with the plasma membrane at the active zone (0 nm from the plasma membrane), that is, there are no lighter pixels between the membranes. Vesicles that were not manually annotated as docked but were 0 nm away from the active zone plasma membrane, were automatically counted as docked when segmentation was quantitated (see below) for data sets counting the number of docked vesicles. Likewise, vesicles annotated as docked were automatically placed in the 0 nm bin of vesicle distances from the plasma membrane. To minimize bias, error, and maintain consistency, all image segmentation, still in the form of randomized files, was thoroughly checked and edited by a second member of the lab. Features were then quantitated using the SynapsEM (**Watanabe et al., 2020**) family of MATLAB (MathWorks) scripts (https://github.com/shigekiwatanabe/SynapsEM). Example electron micrographs shown were adjusted in brightness and contrast to different degrees (depending on the varying brightness and contrast of the raw images), rotated, and cropped in ImageJ before import into Adobe Illustrator.

### In-gel fluorescence assay

SYT7a half-life calculations were determined using data obtained via in-gel fluorescence assays. Rat cortical neurons were transduced with SYT7a-HaloTag expression vectors at 5 DIV. At 13 DIV, neurons were incubated with 100 nM JF635 for 30 minutes at 37°C. These neurons were then washed thre times with conditioned NBMA, the final wash was replaced with conditioned NBMA from sister cultures. Samples were harvested at 13 DIV (post label day 0), 15, 17, 19, and 21 DIV (post label day 2, 4, 6, and 8 respectively) and subjected to standard SDS-PAGE. Gels were analyzed using a BioRad Chemidoc MP imager (BioRad) using far red fluorescence excitation and emission filters. Data were quantified by densitometry using Fiji (**Schindelin et al., 2012**).

### Immunocytochemistry (ICC)

Immunocytochemistry was performed as previously described, (**Vevea & Chapman, 2020**). Primary antibodies were: anti-SYT1 C2A (1:100, 48) (lab stock; mAB 48; RRID:AB_2199314), anti-synaptophysin (SYP) (1:500) (SySy; 101 004; RRID:AB_1210382), anti-SYT7 (1:100) (SySy; 105 173; RRID:AB_887838), anti-pan-neurofascin (1:200) (UC Davis/NIH NeuroMab; 75-172; RRID: AB_2282826), anti-SEC61A (1:100) (Abcam; ab183046; RRID: AB_2620158), anti-GM130 (1:100) (BD Biosciences; 610822; RRID: AB_398141), anti-TGN38/46 (1:20) (abcam; ab2809; RRID: AB_2203290), anti-EEA1 (1:50) (abcam; ab2900; RRID: AB_2262056), anti-M6PR (1:100) (abcam; ab2733; RRID: AB_2122792), anti-sortilin (1:100) (abcam; ab16640; RRID: AB_2192606). Secondary antibodies used include, goat anti-mouse IgG1 IgG-Alexa Fluor 488 (1:500) (Thermofisher; A-21121, RRID:AB_2535764), goat anti-guinea pig IgG-Alexa Fluor 488 (1:500) (Thermofisher; A-11073; RRID:AB_2534117), goat anti-mouse IgG2a-Alexa Fluor 488 (1:500) (Thermofisher; A-21131; RRID:AB_2535771), goat anti-rabbit IgG-Alexa Fluor 488 (1:500) (Thermofisher; A-11008; RRID:AB_143165), goat anti-rabbit IgG-Alexa Fluor 546 (1:500) (Thermofisher; A-11035; RRID:AB_2534093), goat anti-mouse IgG2a-Alexa Fluor 546 (1:500) (Thermofisher; A-21133; RRID:AB_2535772), goat anti-mouse IgG2a-Alexa Fluor 647 (1:500) (Thermofisher; A-21241; RRID:AB_2535810), goat anti-mouse IgG2b-Alexa Fluor 546 (1:500) (Thermofisher; A-21143; RRID:AB_2535779). Images for Figure 3, 4, 5, and 6 were acquired on a Zeiss LSM 880 with a 63x 1.4 NA oil immersion objective using the Airyscan super-resolution detector, and deconvolved using automatic Airyscan settings. For figure 5, identical laser and gain settings were used for conditions in each genotype. The same linear brightness and contrast adjustments were applied to all conditions in their respective genotype.

### Janelia Fluor® dye usage

Haloligand conjugated Janelia Fluor (JF) dyes were graciously provided by Luke Lavis from the Janelia Research Campus. We made use of JF549, JF635, JF646, and JF549i. For normal protein localization in live neurons or HEK293T cells, cultures were incubated with 100 nM JF549, JF635, or JF646, for 30 to 60 min at 37°C. Cultures were washed once and imaged. For ICC experiments that required a JF label, a JF dye was added to the primary antibody mix and incubated overnight at 4°C. For the experiment in Fig. 5e and S5e, we used neurons transduced with HaloTag-SYT7α and incubated with 1 nM of the impermeant JF549i dye for 2 days at 37°C. Incubation for 4 days showed no detectable non-specific uptake of this dye. This dye allowed us to determine if the amino terminal portion of SYT7a transited through the plasma membrane before being cleaved by g-secretase intracellularly.

### Live-cell imaging (excluding iGluSnFR)

HEK293T (Fig. S3g, S4g, and S6a) and primary rodent neuronal (Fig. 3b,d,f and S3a,b,c,e) cultures were transiently transfected with various constructs (pEF and pF(UG) hSyn based) using Lipofectamine™ LTX reagent with PLUS reagent (Thermofisher; A12621) and imaged using the Zeiss LSM880 with Airyscan confocal microscope. Coverslips containing HEK293T or neuronal cultures were placed in standard imaging media (ECF (extracellular fluid/ECF) consisting of 140 mM NaCl, 5 mM KCl, 2 mM CaCl_2_, 2 mM MgCl_2_, 5.5 mM glucose, 20 mM HEPES (pH 7.3), B27 (Gibco), glutamax (Gibco), loaded into the microscope and maintained at physiological temperature (~35°C) and humidity. Cells were imaged in Fast Airyscan mode and processed with automatic Airyscan deconvolution settings after image acquisition. Neurons were incubated with 0.5 mM Prosense 680 (PerkinElmer; NEV10003) overnight (10-12 hrs) to reveal active lysosomes. Incubation for longer time periods, up to 5 days were tested, revealed no obvious cytotoxicity, or improvement in number or fluorescence magnitude of prosense 680 signal.

### iGluSnFR imaging and quantification

Synaptic vesicle exocytosis and glutamate release was monitored via iGluSnFR imaging as previously described (**Vevea & Chapman, 2020**), with the following modifications. Images were acquired at 2×2 binning using the low affinity iGluSnFR (S72A mutation) (**Marvin et al., 2018**), and imaging media contained 2 mM Ca^2+^. For single stimuli imaging, 150 frames were collected at 10 ms exposure (1.5 sec total) and a single field stimulus was triggered at half a second after the initial frame. For paired-pulse imaging, two field stimuli were triggered 50 (20 Hz), 100 (10 Hz), 200 (5 Hz), or 500 ms (2 Hz) apart. As above, 150 frames were collected using a 10 ms exposure (1.5 sec total), with stimuli after 500 ms baseline. For high frequency train stimulation (HFS) 50 stimuli were triggered from field depolarizations at 20 Hz (2.5 sec), and 350 frames with 10 ms exposure were collected (3.5 sec), with HFS start after 500 ms baseline. Glutamate peaks were recorded when the signal was > 4x S.D. of the noise. This was used as the threshold to identify ROIs that released glutamate and for quantification regarding active synapses.

### Colocalization quantification

Pearson’s colocalization was quantified as described previously (**Vevea & Chapman, 2020**), using Fiji for ImageJ and Just Another Colocalization Plugin (JACoP) (**Bolte & Cordelieres, 2006**). Groups were quantified and the simple difference between DAPT and control conditions, with error propagated, is displayed in Figure 6d.

### Compounds and chemicals

Protease inhibitors were DAPT, N-[N-(3,5-Difluorophenacetyl)-L-alanyl]-S-phenylglycine t-butyl Ester, (Apexbio; GSI-IX), GI254023X (MedChemExpress; HY-199956), TAPI-1 (ApexBio; B4686), Verubecestat (ApexBio; MK-8931). The palmitoylation inhibitor was 2-bromopalmitate (Sigma; 238422-10G).

### Statistics

Exact values from experiments and analysis, including number of data points (n) and number of trials for each experiment are listed in the Figure Legends. Electron microscopy data are from 2554 total images (2-D synaptic profiles) from 3 experiments (true biological replicates, different cultures/litters frozen on different days and each imaged and analyzed separately as their own batch). Analysis was done with GraphPad Prism 8.4.3 (GraphPad Software Inc). Data sets were tested for normality using the Anderson-Darling test; if normal, parametric statistical methods were used to analyze data, if not normal, nonparametric methods were used for analysis.

## Supplemental figure legends

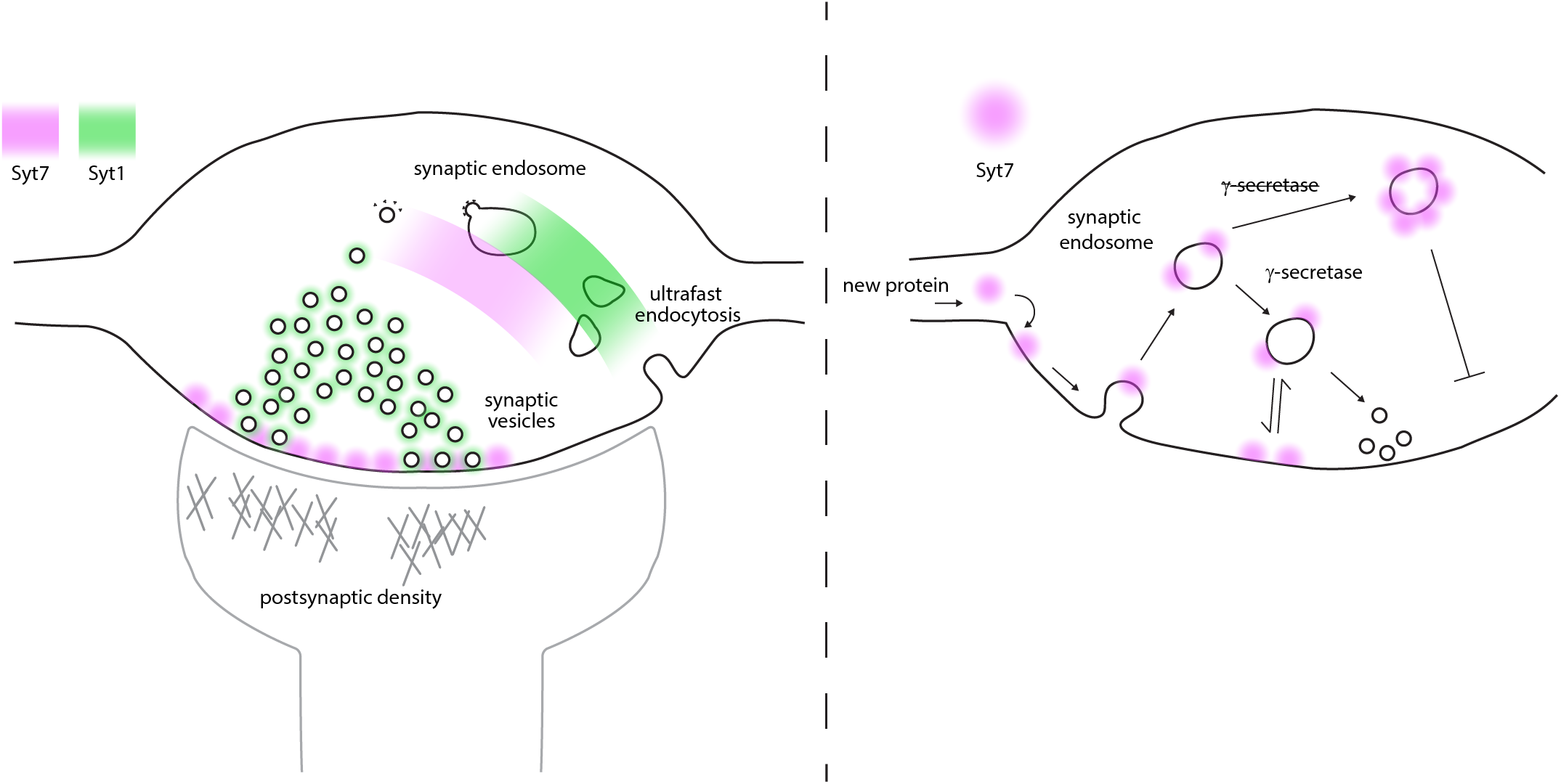
Graphical abstract. **Left**) A model synapse with roles and locations for SYT1 and SYT7 indicated by green and magenta shading in the presynapse area. Historical research on SYT1 has localized SYT1 to the synaptic vesicle (SV) membrane and has supported a role for SYT1 in fast synchronous SV release as well as a role for SYT1 in endocytosis. This work has shown that SYT7 physically and functionally localizes to the plasma membrane of the axon, plays a role in supporting release during short-term synaptic activity, as well as reforming SVs. **Right**) The location of SYT7 (magenta) at the axonal membrane is dependent on γ-secretase processing and palmitoylation near the trans-membrane domain. If palmitoylation is absent (cys -> ala mutations), SYT7 is rapidly degraded. If γ-secretase processing is inhibited, SYT7 mislocalizes to endo-lysosomal intermediate structures.

**Figure S1.**
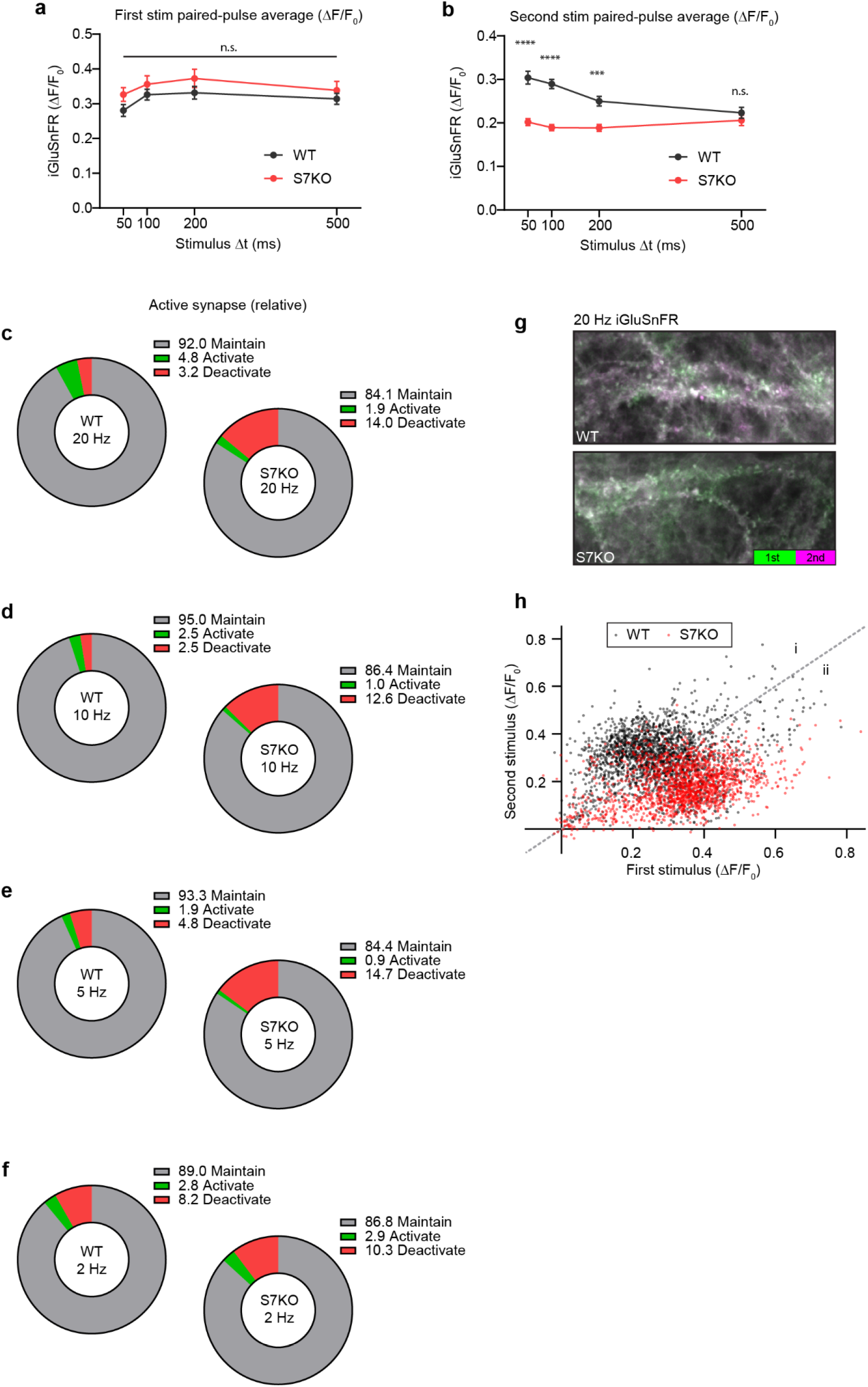
Related to Figure 1. **a**) Quantification of iGluSnFR ΔF/F_0_ peaks from first stimulation during PPR trials. No difference in initial release between any stimulation parameter. Values are means +/− SEM. **b**) Quantification of iGluSnFR ΔF/F_0_ peaks from second stimulation during PPR trials. Values are means +/− SEM. P-values are **** < 0.0001, *** = 0.0004, by two-way ANOVA with Sidak’s multiple comparisons test, full statistics are provided in supplementary statistics table 3. **c-f**) Pie charts showing percentage of ROIs that either maintain glutamate release (grey), activate (green), or deactivate (red) in response to an initial stimulus at the labeled stimulation frequencies. Criteria for glutamate release was based on iGluSnFR ΔF/F_0_ peak amplitude in relation to baseline noise (>4 SD above noise). **g**) Representative images of WT (top) and SYT7KO (bottom) iGluSnFR change in fluorescence in response to two stimulations 50 ms apart (20 Hz). Images are max t-projections of temporally color coded timeseries. Fluorescence from the first stimulus is coded in green (0 – 50 ms post stimulus) and the fluorescence from the second stimulus is coded in purple (50 – 100 ms post stimulus). **h**) Scatterplot from 20 Hz dataset of first stimulus peak (X) by second stimulus peak (Y), WT (black) and SYT7KO (red). Population scatter illustrates difference between WT and SYT7KO ROIs where WT ROIs are grouped predominantly in area (i) because the second stimulus results in facilitation, whereas SYT7KO groups in (ii) because of loss of facilitation.

**Figure S2.**
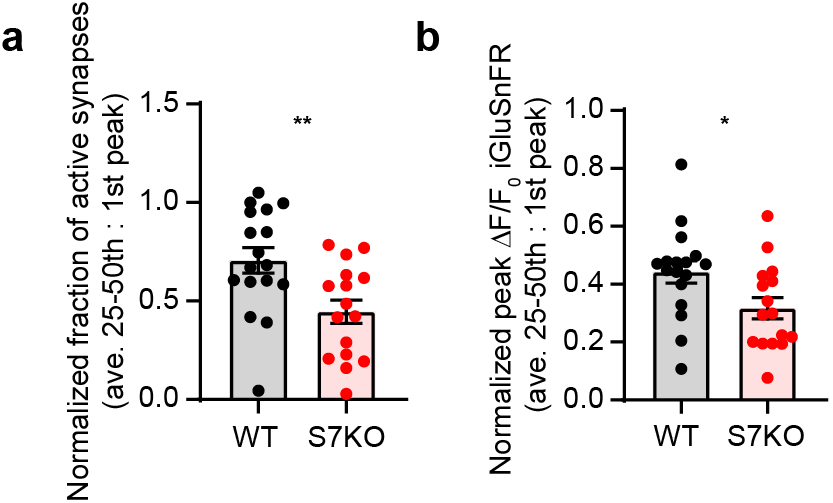
Related to Figure 2. **a**) Quantification of depression as a measure of average active synapses from the 25^th^ to 50^th^ stimulus normalized to the 1^st^ stimulus. Same data set as in Figure 2b-2f. WT (0.71 +/− 0.06) and SYT7KO (0.45 +/− 0.06), values are mean +/− SEM and ** = p-value of 0.0058 using unpaired, two-tailed t-test. **b**) Quantification of depression as a measure of average peak iGluSnFR ΔF/F_0_ from the 25^th^ to 50^th^ stimulus normalized to the 1^st^ stimulus. Same data set as in Figure 2b-2f. WT (0.44 +/− 0.04) and SYT7KO (0.32 +/− 0.04), values are mean +/− SEM and * = p-value of 0.0256 using unpaired, two-tailed t-test.

**Figure S3.**
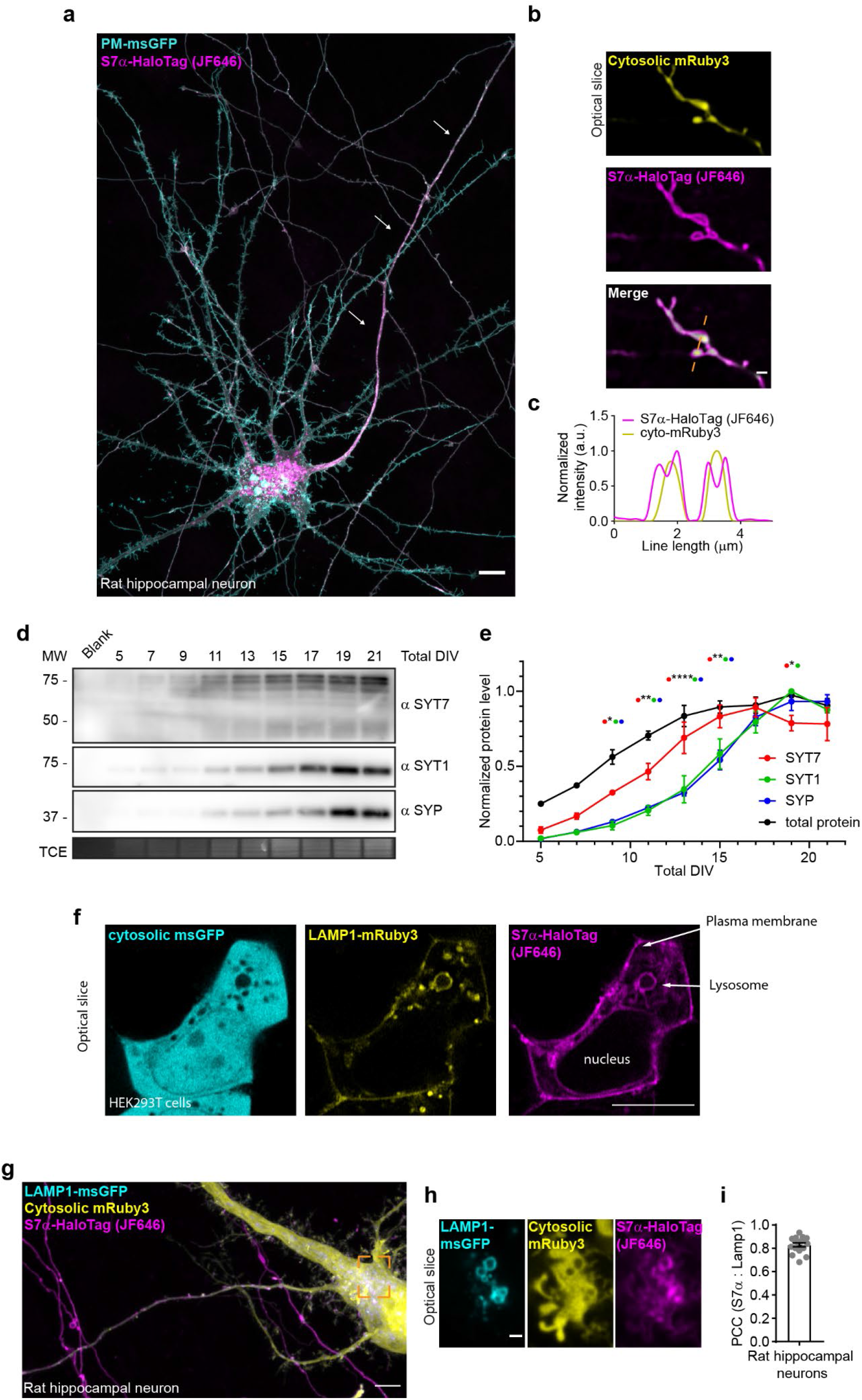
Related to Figure 4. **a**) Representative image of rat hippocampal neuron at 15 DIV transfected with a PM-msGFP P2A SYT7α-HaloTag construct. As with figure 3a) asymmetric localization to the axonal compartment is visible. Scale bar 10 microns. **b**) Additional maximum z-projection of an image of rat hippocampal neurons at 14 DIV expressing transfected LAMP1-msGFP (cyan), cytosolic mRuby3 (yellow), and SYT7α-HaloTag (magenta). Scale bar 5 microns. **c**) A super-resolution optical slice from orange dashed box in b) showing colocalization of LAMP1-msGFP and SYT7α-HaloTag. Scale bar 1 micron. **d**) Bar graph of Pearson’s R statistic (PCC) between SYT7α-HaloTag/JF646 signal and LAMP1-msGFP signal. **e**) Additional super-resolution optical slice of an axon (axonal varicosities) expressing cytosolic mRuby3 (yellow) and SYT7α-HaloTag(magenta). Merged image also denotes line used in f). **f**) graph plotting the normalized intensity profile along the line. Scale bar 1 micron. **g**) Representative super-resolution optical slice of HEK293T cells expressing transfected cytosolic msGFP (cyan), LAMP1-mRuby3 (yellow), and SYT7α-HaloTag (magenta). Scale bar 10 microns. Note in HEK cells, SYT7α also localizes to the plasma membrane and lysosomal membranes. **h**) Representative anti-SYT7, anti-SYT1, anti-Synaptophysin (SYP) immunoblots of developing WT rat cortical neurons at indicated DIV. Trichloroethanol (TCE) is used as a total protein loading control. **i**) Quantification of protein levels normalized to maximal protein signals from immunoblot. Values are mean +/− SEM from 4 independent trials. Significant p-values between SYT7 and SYT1/SYP are labeled, ****<0.0001, ** = 0.001 < x < 0.01, and * = 0.01 < x < 0.05 using two-way ANOVA with Tukey’s correction.

**Figure S4.**
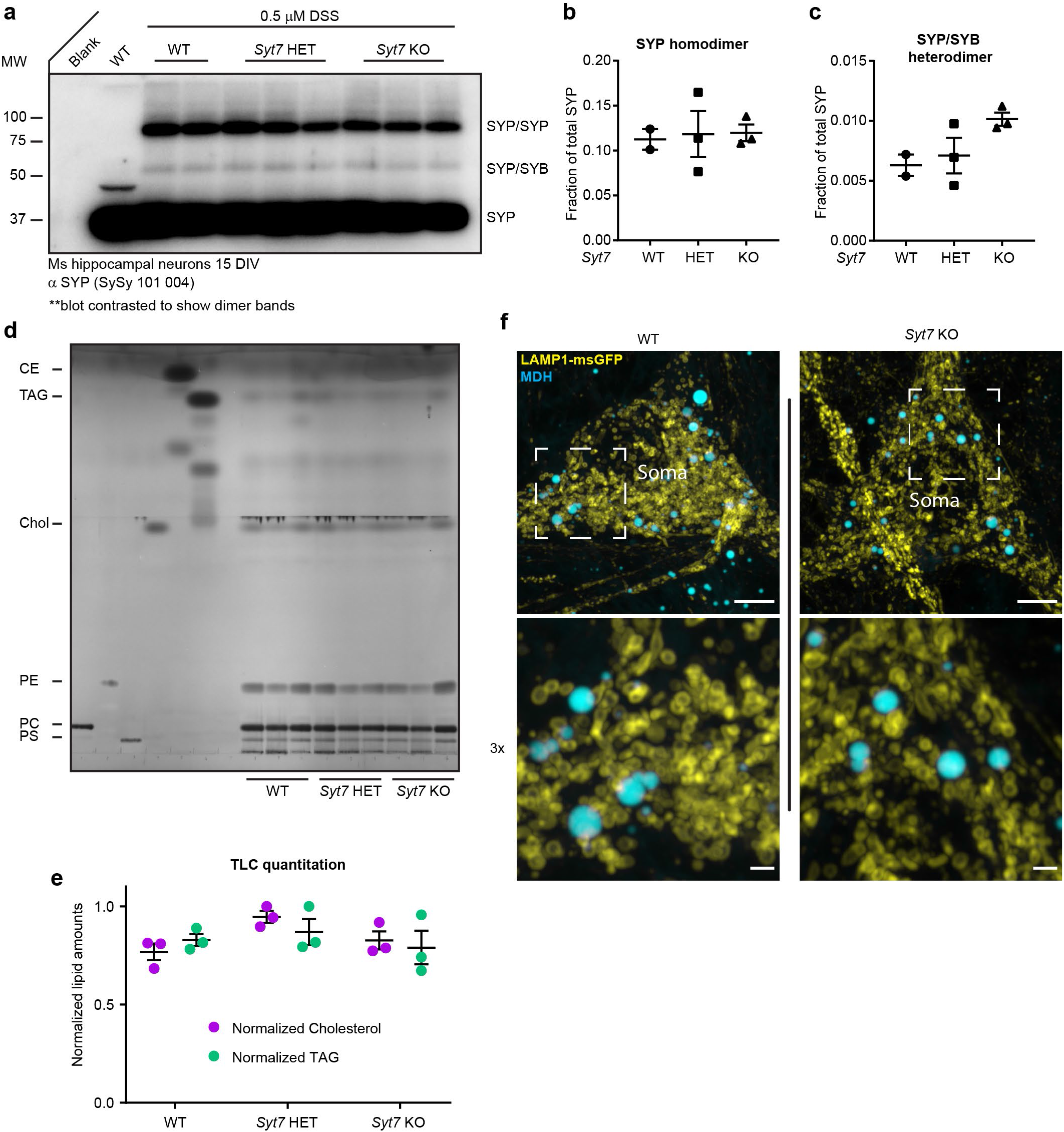
Related to Figure 4. **a**) The anti-SYP immunoblot with WT, *Syt7* HET and *Syt7* KO mouse hippocampal neurons. All samples used in analysis are shown in this blot (2x WT, 3x HET, and 3x KO), a WT sample without DSS treatment is shown as a control. All other samples were incubated in 0.5 mM DSS for 45 min to link physically interacting proteins. **b**) SYP homodimer quantification using densitometry. **c**) SYP/SYB heterodimer quantification using densitometry. **d**) Thin layer chromatography (TLC) examining total lipid levels from mature neurons, phosphatidylserine (PS), phosphatidylcholine (PC), phosphatidylethanolamine (PE), cholesterol (chol), triacylglycerol (TAG), and cholesterol ester (CE). **e**) Quantification of chol and TAG using densitometry. **f**) Representative super-resolution fluorescent images of WT and SYT7KO mouse hippocampal neurons at 15 DIV expressing uniformly transduced LAMP1-msGFP (yellow) and incubated with 50 μM monodansylpentane (MDH) (cyan) for 30 minutes. MDH is a solvochromatic dye that fluoresces in the blue spectrum when it is in a nonpolar environment. It is an excellent dye for lipid droplet labelling (**Vevea & Chapman, 2020**). Scale bar 5 microns top image row and 1 micron bottom (3x) image row.

**Figure S5.**
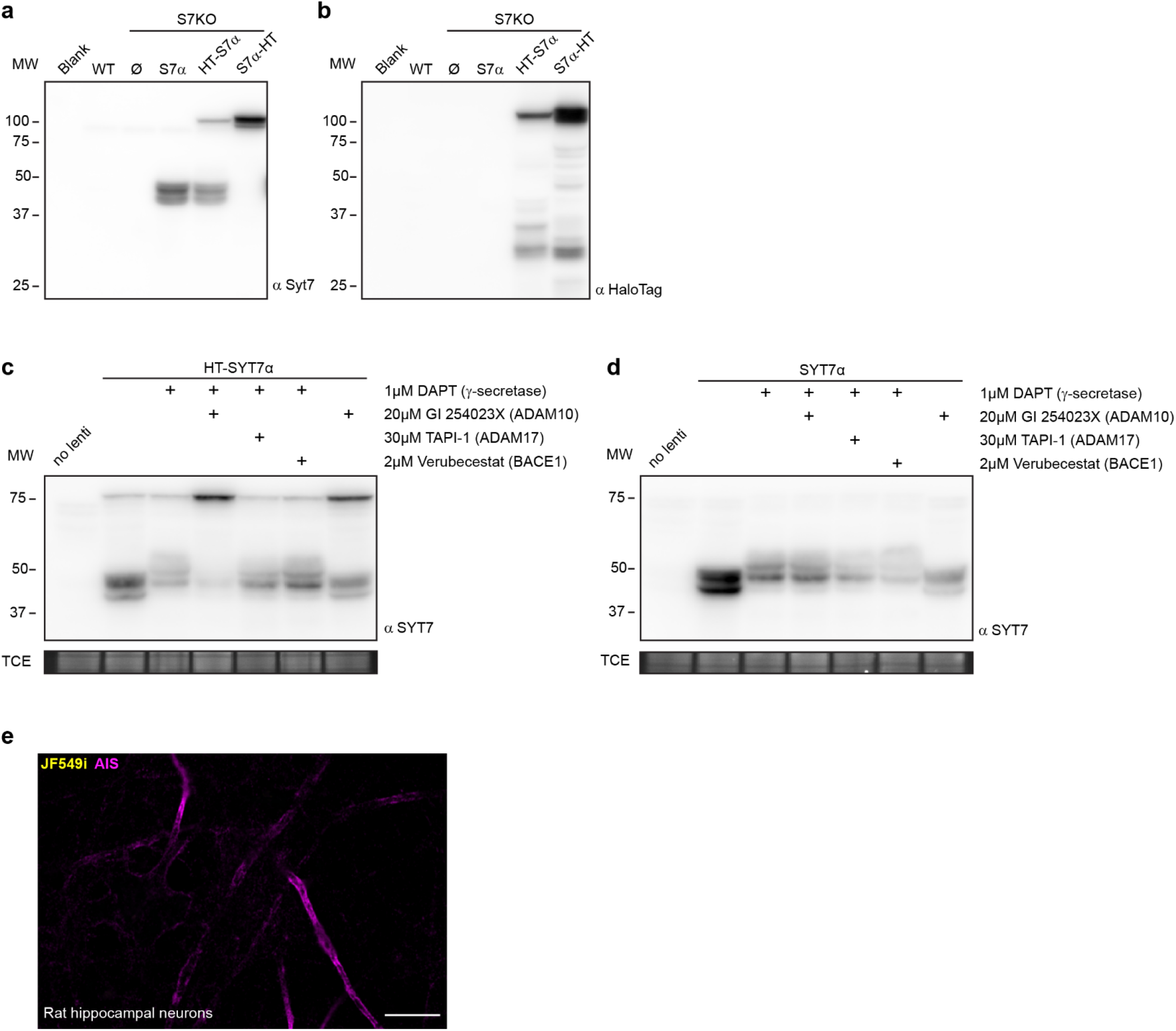
Related to Figure 5. **a**) Representative anti-SYT7 immunoblot from WT and SYT7KO dissociated mouse hippocampal neurons transduced with various constructs. WT (endogenous SYT7 is expressed at very low levels in neurons obtained from P0 mice) and SYT7KO conditions shown as controls. SYT7α represents over-expressed untagged SYT7 alpha isoform, HT-SYT7α and SYT7α-HT represent amino and carboxy terminal tagged SYT7 with the HaloTag enzyme. **b**) The same blot as in a) but stripped and probed with anti-HaloTag. **c**) Representative anti-SYT7 immunoblot of rat cortical neurons expressing transduced HT-SYT7α with various combinations of a competitive γ-secretase inhibitor (DAPT) and metalloprotease inhibitors, GI 254023X for ADAM10, TAPI-1 for ADAM17, and Verubecestat for BACE1. TCE as a load control. **d**) The same setup as in c) but with neurons transduced with untagged SYT7α. TCE as a load control. **e**) Representative super-resolution optical slice of an untransduced rat hippocampal neuron. Before fixing neurons, they were incubated with 1 nM JF549i for 2 days to reveal any nonspecific labelling from the JF549i dye. Fixed neurons were decorated with anti-pan-neurofascin antibodies to mark axon initial segments. **f**) Diagram of proposed SYT7 sensor. Includes preprolactin signal sequence (PP) – HaloTag enzyme – SYT7α TMD (1-87 a.a.) – msGFP – Streptavidin binding protein (SBP). **g**) Representative super-resolution max z-projections of HEK293T cells transfected with construct described in f) under control and inhibitor (+DAPT and GI 254023X) conditions. Scale bar 10 microns.

**Figure S6.**
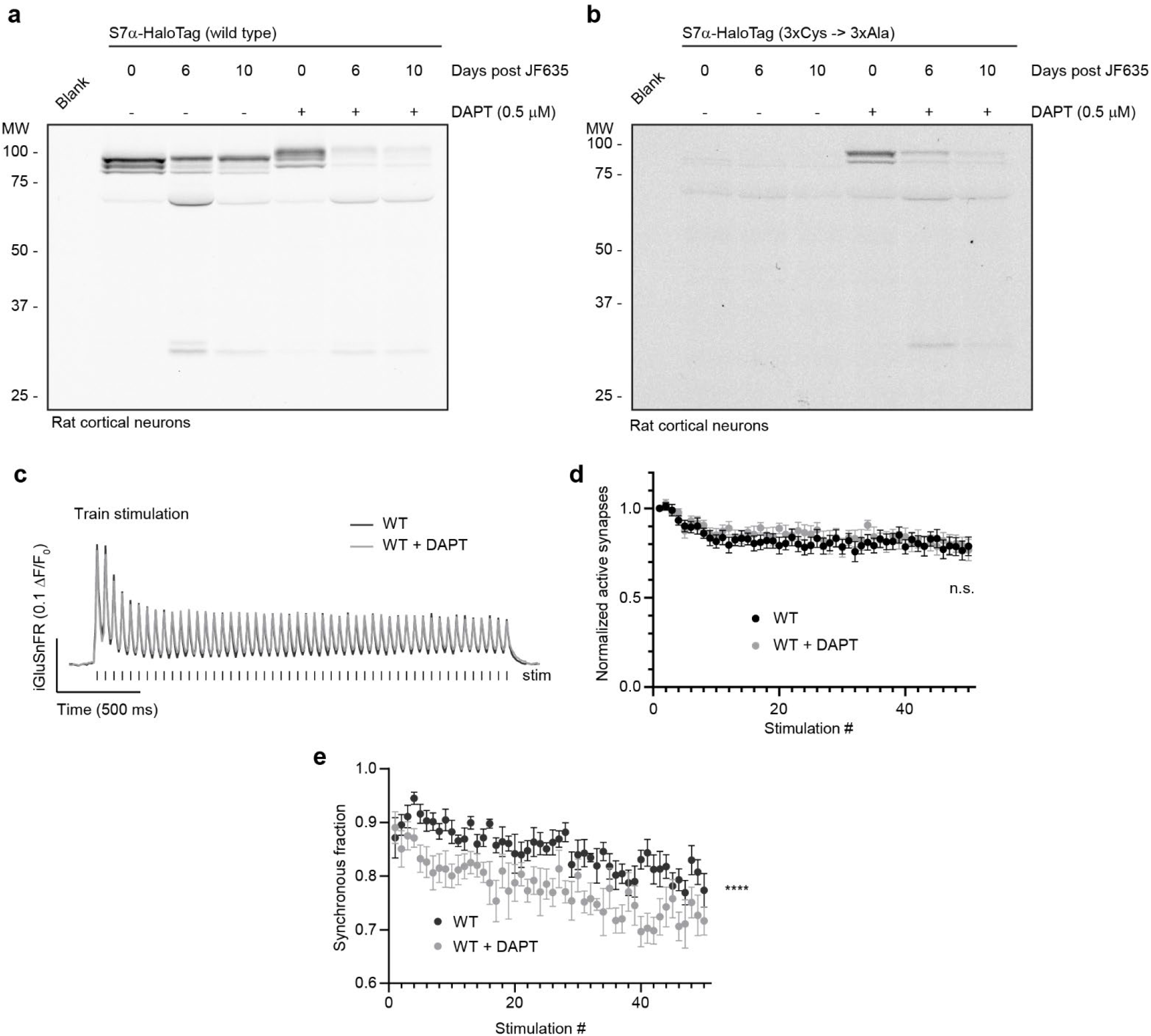
Related to Figure 6. **a**) Representative in-gel fluorescence of protein extracted from rat cortical neurons transduced with SYT7α-HaloTag and pulse-chased with JF635 at 13 DIV under control conditions and when grown in 0.5 μM DAPT. **b**) Representative in-gel fluorescence of protein extracted from rat cortical neurons transduced with palmitylation site mutant (3x Cys -> 3x Ala) SYT7α-HaloTag and pulse-chased with JF635 at 13 DIV under control conditions and when grown in 0.5 μM DAPT. Palmitoylation mutants are not stable in control conditions, presumably because it is cleaved by γ-secretase and degraded (because lack of membrane association from palmitoylation sites). **c**) Average iGluSnFR ΔF/F_0_ traces from HFS of WT (black, n = 9) and WT + 0.5 μM DAPT (grey, n = 11) treated mouse hippocampal neurons, from 2 independent experiments. Samples were field stimulated with a frequency of 20 Hz for 2.5 s (50 APs). **d**) Fraction of active synapses (synapses releasing peak glutamate above baseline, >4 SD above noise) as a function of stimulation number during HFS. Values are means +/− SEM, p = n.s. by two-way ANOVA comparing genotype. **e**) Synchronous fraction of iGluSnFR ΔF/F_0_ peaks (synchronous peaks within 10 ms of stimulus) as a function of stimulation number during HFS. Values are means +/− SEM, p < 0.0001 by two-way ANOVA comparing genotype.

**Figure S7.**
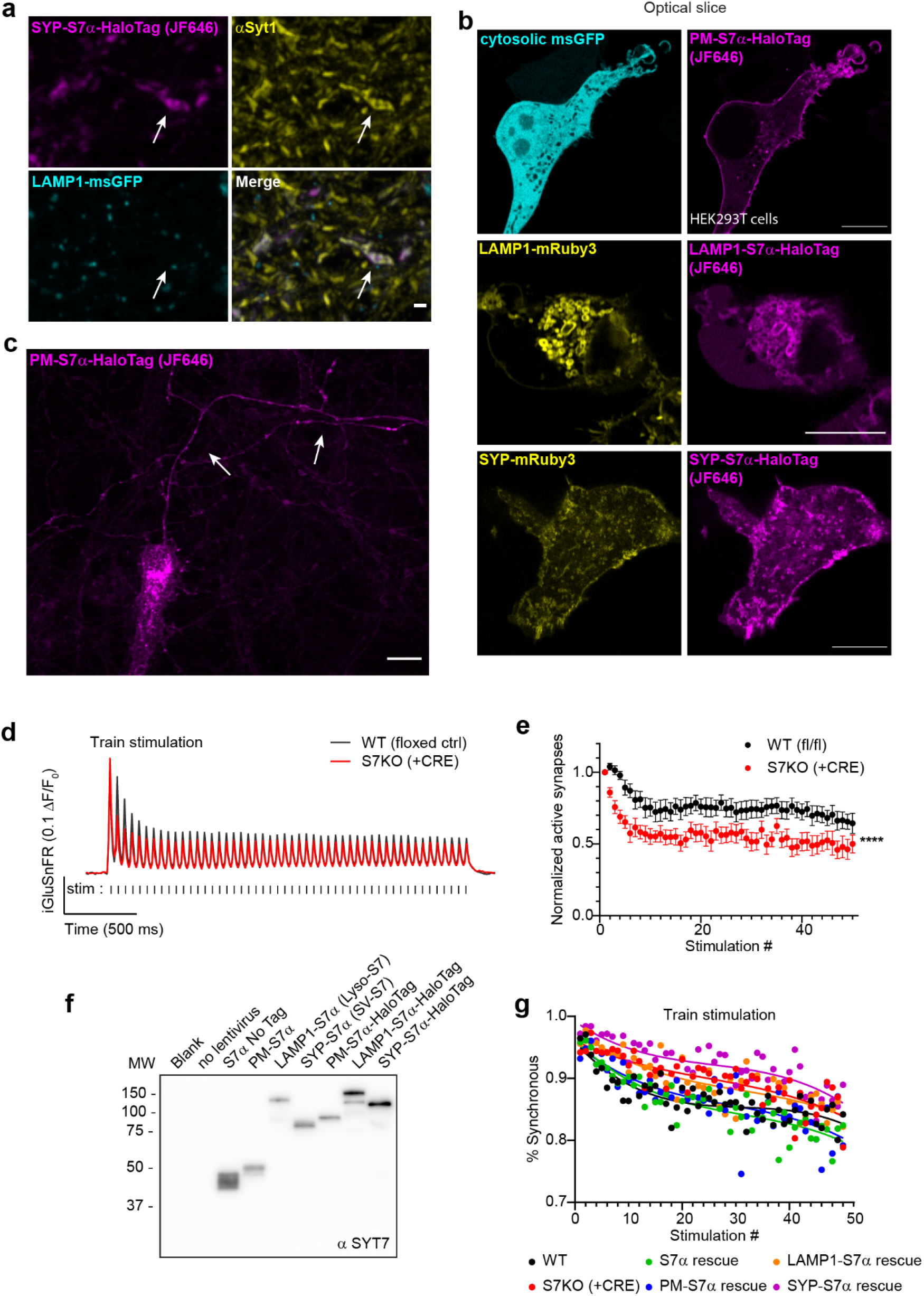
Related to Figure 7. **a**) Representative image taken with the same imaging conditions as in Fig. 7c. Neurons transduced with SYP-SYT7α-HaloTag. Arrows point to a SV cluster and nearby LAMP1+ structure. Scale bar = 1 micron. **b**) Representative super-resolution optical slices of HEK293T cells expressing the location specific SYT7α rescue constructs. From left to right, cytosolic msGFP (cyan) with PM-SYT7α (magenta), LAMP1-mRuby3 (yellow) with LAMP1-SYT7α (magenta), and SYP-mRuby3 (yellow) with SYP-SYT7α (magenta). Scale bar 10 microns. **c**) Same image as shown in Fig. 7a, but shown here with a larger field of view and only the PM-S7α-HaloTag channel. Arrows point to the axon. Scale bar = 10 microns. **d**) Average iGluSnFR ΔF/F_0_ traces from SYT7 floxed strain, WT (floxed/no CRE, black, n = 13) and SYT7KO (+CRE, red, n = 13), from 3 independent experiments. **e**) Fraction of active synapses (synapses releasing peak glutamate above baseline, >4 SD above noise) as a function of stimulation number during HFS. Values are means +/− SEM, p < 0.0001 by two-way ANOVA comparing genotype. **f**) Representative anti-SYT7 immunoblot from rat cortical neurons, illustrating expression of labeled SYT7α constructs used in Fig. 6. **g**) Average synchronous release percentage plotted by stimulation number, genotypes as labeled. As HFS continues, synchronous release decreases. Same data as in Fig. 6h. Lines are non-linear third order polynomial fits to data, for illustrative purposes only.

## Supplemental summary statistics

**supplementary statistics table 1.** For Figure 1e.

| Compare each cell mean with the other cell mean in that row |  |  |  |  |  |  |  |
| --- | --- | --- | --- | --- | --- | --- | --- |
| Number of families | 1 |  |  |  |  |  |  |
| Number of comparisons per family | 4 |  |  |  |  |  |  |
| Alpha | 0.05 |  |  |  |  |  |  |
| Sidak's multiple comparisons test | Predicted (LS) mean diff, | 95,00% CI of diff, | Significant? | Summary | Adjusted P Value |  |  |
| WT - S7KO |  |  |  |  |  |  |  |
| 20 Hz | 0.5768 | 0,3901 to 0,7634 | Yes | **** | <0,0001 |  |  |
| 10 Hz | 0.4322 | 0,2387 to 0,6256 | Yes | **** | <0,0001 |  |  |
| 5 Hz | 0.2763 | 0,08962 to 0,4629 | Yes | ** | 0.0012 |  |  |
| 2 Hz | 0.1031 | -0,09391 to 0,3001 | No | ns | 0.5643 |  |  |
| Test details | Predicted (LS) mean 1 | Predicted (LS) mean 2 | Predicted (LS) mean diff, | SE of diff, | N1 | N2 | t |
| WT - S7KO |  |  |  |  |  |  |  |
| 20 Hz | 1.153 | 0.5767 | 0.5768 | 0.07362 | 14 | 15 | 7.835 |
| 10 Hz | 0.9148 | 0.4827 | 0.4322 | 0.0763 | 14 | 13 | 5.664 |
| 5 Hz | 0.7609 | 0.4846 | 0.2763 | 0.07362 | 15 | 14 | 3.753 |
| 2 Hz | 0.7013 | 0.5982 | 0.1031 | 0.0777 | 13 | 13 | 1.327 |
|  | delta T (ms) | WT |  |  | S7KO |  |  |
|  | X | Mean | SEM | N | Mean | SEM | N |
| 20 Hz | 50 | 1.153452658 | 0.092200939 | 14 | 0.576676482 | 0.029094085 | 15 |
| 10 Hz | 100 | 0.914810555 | 0.069095618 | 14 | 0.482658414 | 0.027627979 | 13 |
| 5 Hz | 200 | 0.760889287 | 0.045842498 | 15 | 0.484637944 | 0.038067777 | 14 |
| 2 Hz | 500 | 0.701312358 | 0.039677713 | 13 | 0.598230091 | 0.049256498 | 13 |

**supplementary statistics table 2.**
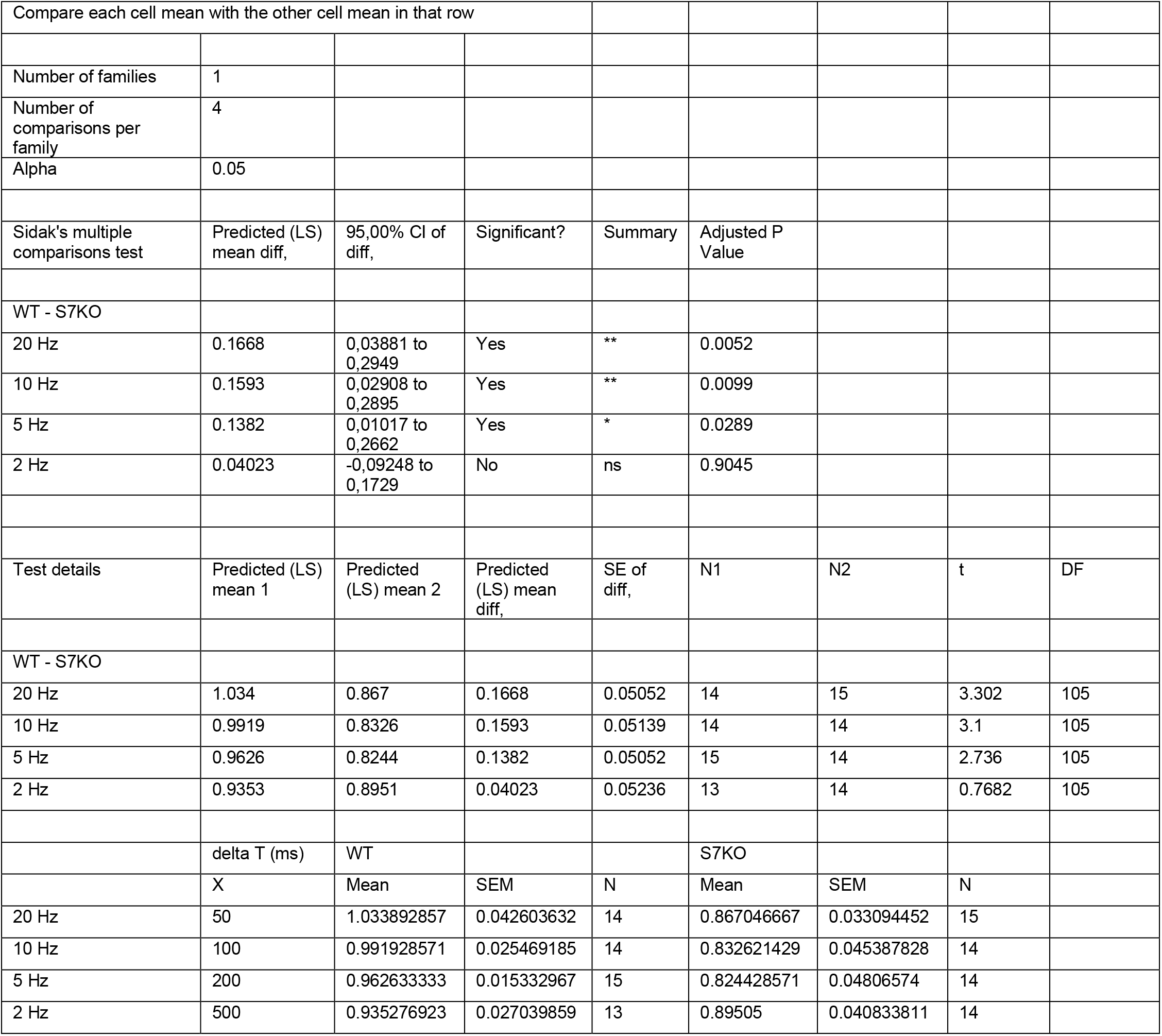
For Figure 1f.

**supplementary statistics table 3.**
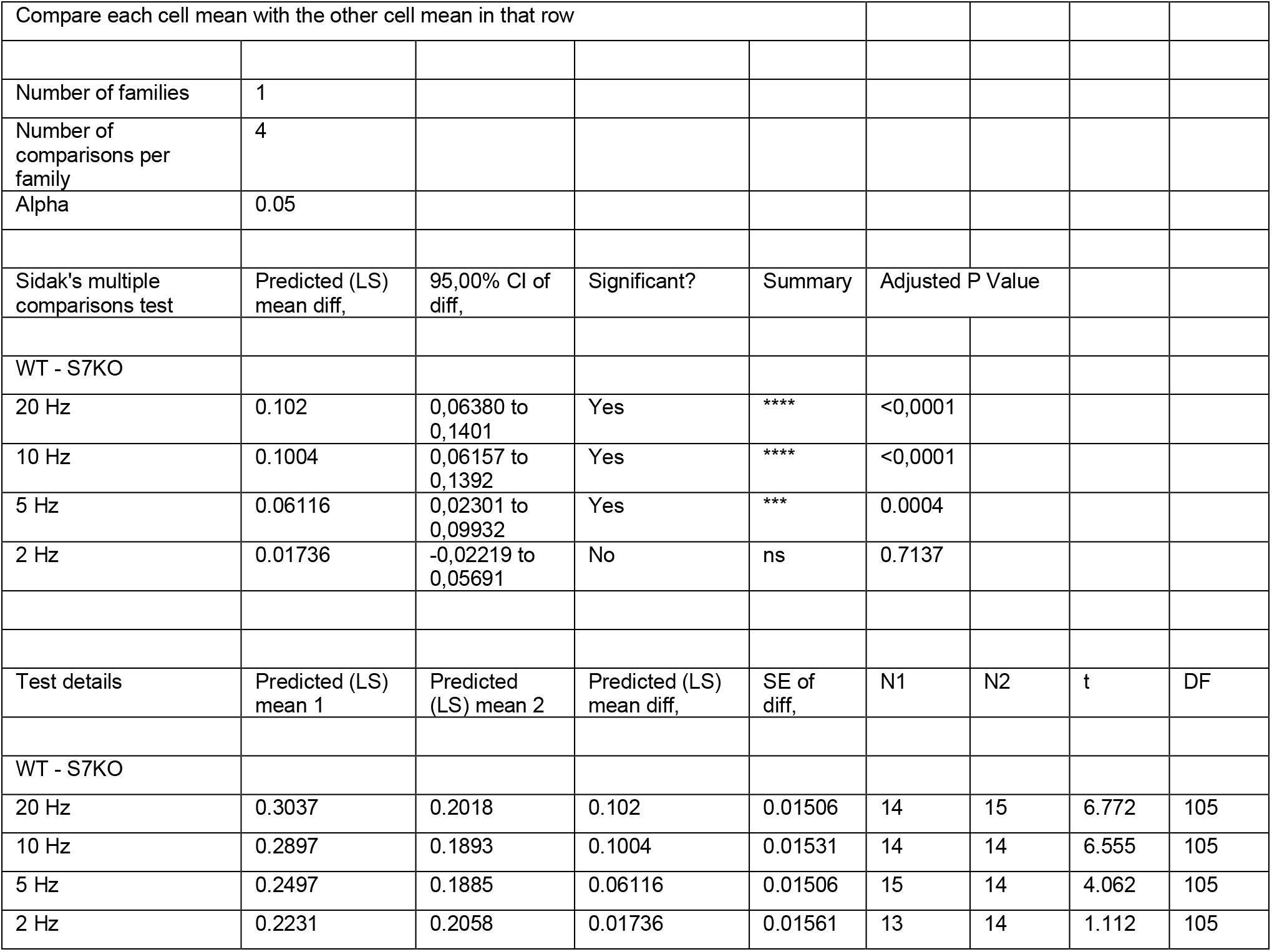
For Figure S1b.

**supplementary statistics table 4.**
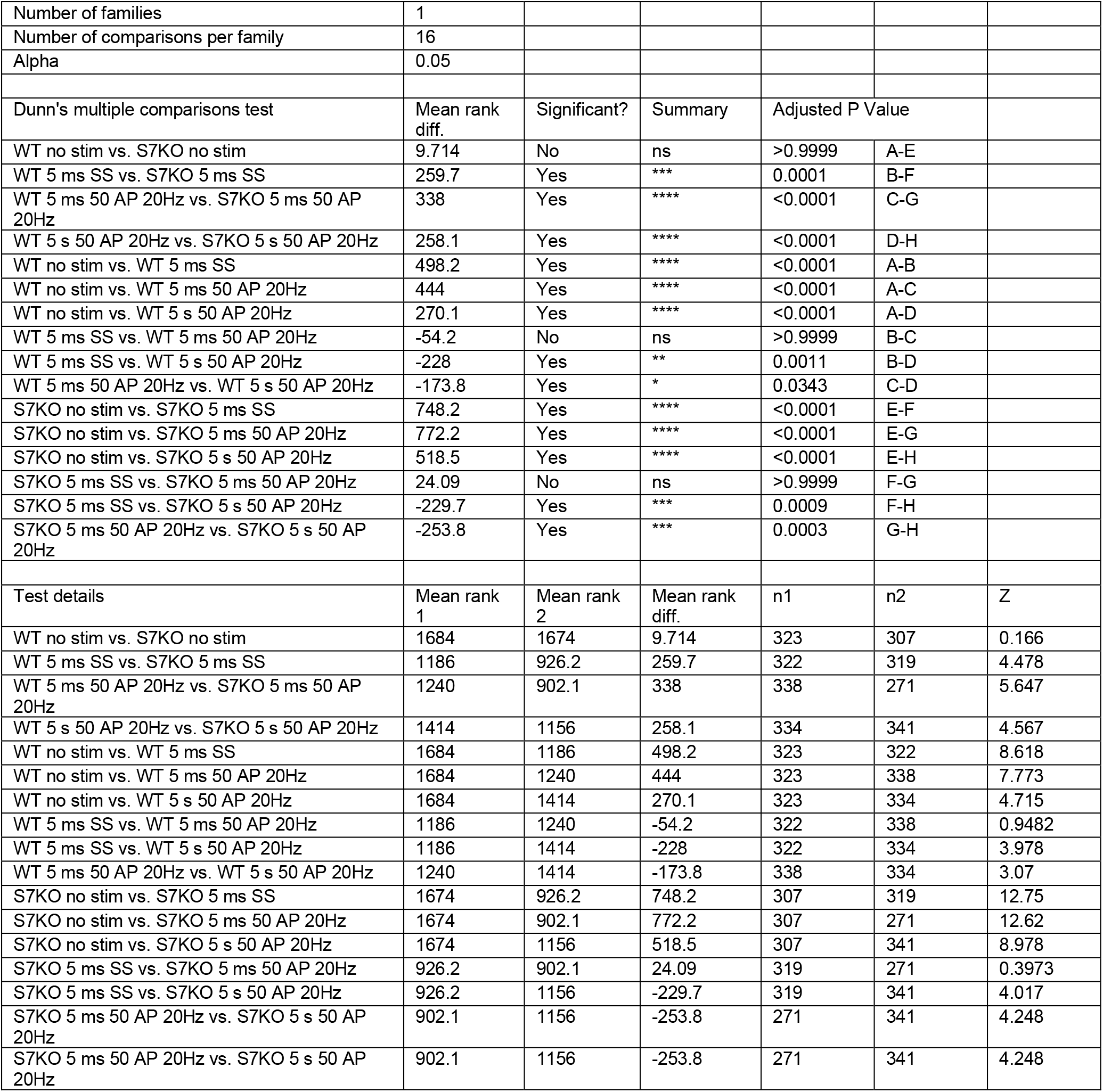
For Figure 3c.

|  |  |  |  |  |  |  |
| --- | --- | --- | --- | --- | --- | --- |
| Number of families | 1 |  |  |  |  |  |
| Number of comparisons per family | 16 |  |  |  |  |  |
| Alpha | 0.05 |  |  |  |  |  |
| Dunn's multiple comparisons test | Mean rank diff. | Significant? | Summary | Adjusted P Value |  |  |
| WT no stim vs. S7KO no stim | 9.714 | No | ns | >0.9999 | A-E |  |
| WT 5 ms SS vs. S7KO 5 ms SS | 259.7 | Yes | *** | 0.0001 | B-F |  |
| WT 5 ms 50 AP 20Hz vs. S7KO 5 ms 50 AP 20Hz | 338 | Yes | **** | <0.0001 | C-G |  |
| WT 5 s 50 AP 20Hz vs. S7KO 5 s 50 AP 20Hz | 258.1 | Yes | **** | <0.0001 | D-H |  |
| WT no stim vs. WT 5 ms SS | 498.2 | Yes | **** | <0.0001 | A-B |  |
| WT no stim vs. WT 5 ms 50 AP 20Hz | 444 | Yes | **** | <0.0001 | A-C |  |
| WT no stim vs. WT 5 s 50 AP 20Hz | 270.1 | Yes | **** | <0.0001 | A-D |  |
| WT 5 ms SS vs. WT 5 ms 50 AP 20Hz | -54.2 | No | ns | >0.9999 | B-C |  |
| WT 5 ms SS vs. WT 5 s 50 AP 20Hz | -228 | Yes | ** | 0.0011 | B-D |  |
| WT 5 ms 50 AP 20Hz vs. WT 5 s 50 AP 20Hz | -173.8 | Yes | * | 0.0343 | C-D |  |
| S7KO no stim vs. S7KO 5 ms SS | 748.2 | Yes | **** | <0.0001 | E-F |  |
| S7KO no stim vs. S7KO 5 ms 50 AP 20Hz | 772.2 | Yes | **** | <0.0001 | E-G |  |
| S7KO no stim vs. S7KO 5 s 50 AP 20Hz | 518.5 | Yes | **** | <0.0001 | E-H |  |
| S7KO 5 ms SS vs. S7KO 5 ms 50 AP 20Hz | 24.09 | No | ns | >0.9999 | F-G |  |
| S7KO 5 ms SS vs. S7KO 5 s 50 AP 20Hz | -229.7 | Yes | *** | 0.0009 | F-H |  |
| S7KO 5 ms 50 AP 20Hz vs. S7KO 5 s 50 AP 20Hz | -253.8 | Yes | *** | 0.0003 | G-H |  |
| Test details | Mean rank 1 | Mean rank 2 | Mean rank diff. | n1 | n2 | Z |
| WT no stim vs. S7KO no stim | 1684 | 1674 | 9.714 | 323 | 307 | 0.166 |
| WT 5 ms SS vs. S7KO 5 ms SS | 1186 | 926.2 | 259.7 | 322 | 319 | 4.478 |
| WT 5 ms 50 AP 20Hz vs. S7KO 5 ms 50 AP 20Hz | 1240 | 902.1 | 338 | 338 | 271 | 5.647 |
| WT 5 s 50 AP 20Hz vs. S7KO 5 s 50 AP 20Hz | 1414 | 1156 | 258.1 | 334 | 341 | 4.567 |
| WT no stim vs. WT 5 ms SS | 1684 | 1186 | 498.2 | 323 | 322 | 8.618 |
| WT no stim vs. WT 5 ms 50 AP 20Hz | 1684 | 1240 | 444 | 323 | 338 | 7.773 |
| WT no stim vs. WT 5 s 50 AP 20Hz | 1684 | 1414 | 270.1 | 323 | 334 | 4.715 |
| WT 5 ms SS vs. WT 5 ms 50 AP 20Hz | 1186 | 1240 | -54.2 | 322 | 338 | 0.9482 |
| WT 5 ms SS vs. WT 5 s 50 AP 20Hz | 1186 | 1414 | -228 | 322 | 334 | 3.978 |
| WT 5 ms 50 AP 20Hz vs. WT 5 s 50 AP 20Hz | 1240 | 1414 | -173.8 | 338 | 334 | 3.07 |
| S7KO no stim vs. S7KO 5 ms SS | 1674 | 926.2 | 748.2 | 307 | 319 | 12.75 |
| S7KO no stim vs. S7KO 5 ms 50 AP 20Hz | 1674 | 902.1 | 772.2 | 307 | 271 | 12.62 |
| S7KO no stim vs. S7KO 5 s 50 AP 20Hz | 1674 | 1156 | 518.5 | 307 | 341 | 8.978 |
| S7KO 5 ms SS vs. S7KO 5 ms 50 AP 20Hz | 926.2 | 902.1 | 24.09 | 319 | 271 | 0.3973 |
| S7KO 5 ms SS vs. S7KO 5 s 50 AP 20Hz | 926.2 | 1156 | -229.7 | 319 | 341 | 4.017 |
| S7KO 5 ms 50 AP 20Hz vs. S7KO 5 s 50 AP 20Hz | 902.1 | 1156 | -253.8 | 271 | 341 | 4.248 |
| S7KO 5 ms 50 AP 20Hz vs. S7KO 5 s 50 AP 20Hz | 902.1 | 1156 | -253.8 | 271 | 341 | 4.248 |

**supplementary statistics table 5.**
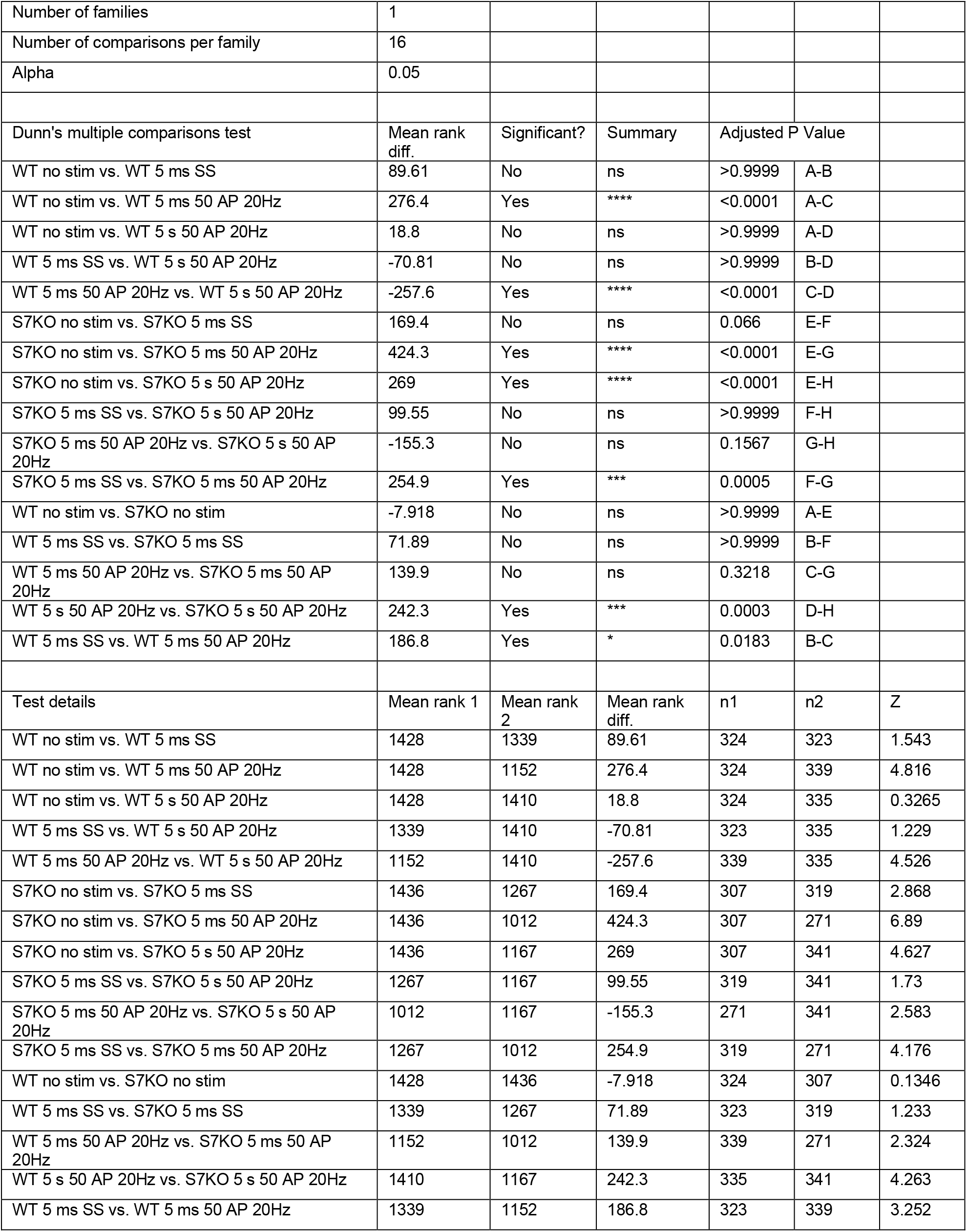
For Figure 3d.

**supplementary statistics table 6.** For Figure 7f.

| Compare column means (main column effect) |  |  |  |  |  |  |  |  |
| --- | --- | --- | --- | --- | --- | --- | --- | --- |
| Number of families | 1 |  |  |  |  |  |  |  |
| Number of comparisons per family | 15 |  |  |  |  |  |  |  |
| Alpha | 0.05 |  |  |  |  |  |  |  |
| Tukey's multiple comparisons test | Predicted (LS) mean diff, | 95,00% CI of diff, | Significant? | Summary | Adjusted P Value |  |  |  |
| WT vs. S7KO (+CRE) | 0.2344 | 0,1995 to 0,2694 | Yes | **** | <0,0001 |  |  |  |
| WT vs. S7a Rescue | 0.07192 | 0,03824 to 0,1056 | Yes | **** | <0,0001 |  |  |  |
| WT vs. PM-S7a Rescue | 0.08996 | 0,05628 to 0,1236 | Yes | **** | <0,0001 |  |  |  |
| WT vs. Lyso-S7a Rescue | 0.05774 | 0,02405 to 0,09142 | Yes | **** | <0,0001 |  |  |  |
| WT vs. SV-S7a Rescue | 0.2143 | 0,1811 to 0,2474 | Yes | **** | <0,0001 |  |  |  |
| S7KO (+CRE) vs. S7a Rescue | -0.1625 | -0,1975 to -0,1275 | Yes | **** | <0,0001 |  |  |  |
| S7KO (+CRE) vs. PM-S7a Rescue | -0.1445 | -0,1794 to -0,1095 | Yes | **** | <0,0001 |  |  |  |
| S7KO (+CRE) vs. Lyso-S7a Rescue | -0.1767 | -0,2116 to -0,1417 | Yes | **** | <0,0001 |  |  |  |
| S7KO (+CRE) vs. SV-S7a Rescue | -0.02014 | -0,05458 to 0,01431 | No | ns | 0.554 |  |  |  |
| S7a Rescue vs. PM-S7a Rescue | 0.01804 | -0,01564 to 0,05172 | No | ns | 0.6467 |  |  |  |
| S7a Rescue vs. Lyso-S7a Rescue | -0.01418 | -0,04786 to 0,01950 | No | ns | 0.8368 |  |  |  |
| S7a Rescue vs. SV-S7a Rescue | 0.1424 | 0,1092 to 0,1755 | Yes | **** | <0,0001 |  |  |  |
| PM-S7a Rescue vs. Lyso-S7a Rescue | -0.03222 | -0,06590 to 0,001460 | No | ns | 0.0701 |  |  |  |
| PM-S7a Rescue vs. SV-S7a Rescue | 0.1243 | 0,09118 to 0,1575 | Yes | **** | <0,0001 |  |  |  |
| Lyso-S7a Rescue vs. SV-S7a Rescue | 0.1565 | 0,1234 to 0,1897 | Yes | **** | <0,0001 |  |  |  |
| Test details | Predicted (LS) mean 1 | Predicted (LS) mean 2 | Predicted (LS) mean diff, | SE of diff, | N1 | N2 | q | DF |
| WT vs. S7KO (+CRE) | 0.8053 | 0.5708 | 0.2344 | 0.01226 | 750 | 650 | 27.04 | 4150 |
| WT vs. S7a Rescue | 0.8053 | 0.7333 | 0.07192 | 0.01181 | 750 | 750 | 8.609 | 4150 |
| WT vs. PM-S7a Rescue | 0.8053 | 0.7153 | 0.08996 | 0.01181 | 750 | 750 | 10.77 | 4150 |
| WT vs. Lyso-S7a Rescue | 0.8053 | 0.7475 | 0.05774 | 0.01181 | 750 | 750 | 6.911 | 4150 |
| WT vs. SV-S7a Rescue | 0.8053 | 0.591 | 0.2143 | 0.01163 | 750 | 800 | 26.06 | 4150 |
| S7KO (+CRE) vs. S7a Rescue | 0.5708 | 0.7333 | -0.1625 | 0.01226 | 650 | 750 | 18.75 | 4150 |
| S7KO (+CRE) vs. PM-S7a Rescue | 0.5708 | 0.7153 | -0.1445 | 0.01226 | 650 | 750 | 16.66 | 4150 |
| S7KO (+CRE) vs. Lyso-S7a Rescue | 0.5708 | 0.7475 | -0.1767 | 0.01226 | 650 | 750 | 20.38 | 4150 |
| S7KO (+CRE) vs. SV-S7a Rescue | 0.5708 | 0.591 | -0.02014 | 0.01208 | 650 | 800 | 2.357 | 4150 |
| S7a Rescue vs. PM-S7a Rescue | 0.7333 | 0.7153 | 0.01804 | 0.01181 | 750 | 750 | 2.16 | 4150 |
| S7a Rescue vs. Lyso-S7a Rescue | 0.7333 | 0.7475 | -0.01418 | 0.01181 | 750 | 750 | 1.698 | 4150 |
| S7a Rescue vs. SV-S7a Rescue | 0.7333 | 0.591 | 0.1424 | 0.01163 | 750 | 800 | 17.32 | 4150 |
| PM-S7a Rescue vs. Lyso-S7a Rescue | 0.7153 | 0.7475 | -0.03222 | 0.01181 | 750 | 750 | 3.857 | 4150 |
| PM-S7a Rescue vs. SV-S7a Rescue | 0.7153 | 0.591 | 0.1243 | 0.01163 | 750 | 800 | 15.12 | 4150 |
| Lyso-S7a Rescue vs. SV-S7a Rescue | 0.7475 | 0.591 | 0.1565 | 0.01163 | 750 | 800 | 19.04 | 4150 |

**supplementary statistics table 7.** For Figure 7h.

| Compare column means (main column effect) |  |  |  |  |  |  |  |  |
| --- | --- | --- | --- | --- | --- | --- | --- | --- |
| Number of families | 1 |  |  |  |  |  |  |  |
| Number of comparisons per family | 15 |  |  |  |  |  |  |  |
| Alpha | 0.05 |  |  |  |  |  |  |  |
| Sidak's multiple comparisons test | Predicted (LS) mean diff, | 95,00% CI of diff, | Significant? | Summary | Adjusted P Value |  |  |  |
| WT vs. S7KO (+CRE) | -0.03159 | -0,04868 to -0,01451 | Yes | **** | <0,0001 |  |  |  |
| WT vs. S7a Rescue | 0.007326 | -0,009140 to 0,02379 | No | ns | 0.9595 |  |  |  |
| WT vs. PM-S7a Rescue | -0.0004883 | -0,01697 to 0,01599 | No | ns | >0,9999 |  |  |  |
| WT vs. Lyso-S7a Rescue | -0.02815 | -0,04462 to -0,01169 | Yes | **** | <0,0001 |  |  |  |
| WT vs. SV-S7a Rescue | -0.05557 | -0,07177 to -0,03936 | Yes | **** | <0,0001 |  |  |  |
| S7KO (+CRE) vs. S7a Rescue | 0.03892 | 0,02183 to 0,05601 | Yes | **** | <0,0001 |  |  |  |
| S7KO (+CRE) vs. PM-S7a Rescue | 0.0311 | 0,01401 to 0,04820 | Yes | **** | <0,0001 |  |  |  |
| S7KO (+CRE) vs. Lyso-S7a Rescue | 0.003441 | -0,01365 to 0,02053 | No | ns | >0,9999 |  |  |  |
| S7KO (+CRE) vs. SV-S7a Rescue | -0.02397 | -0,04081 to -0,007134 | Yes | *** | 0.0005 |  |  |  |
| S7a Rescue vs. PM-S7a Rescue | -0.007814 | -0,02429 to 0,008664 | No | ns | 0.9329 |  |  |  |
| S7a Rescue vs. Lyso-S7a Rescue | -0.03548 | -0,05194 to -0,01901 | Yes | **** | <0,0001 |  |  |  |
| S7a Rescue vs. SV-S7a Rescue | -0.06289 | -0,07910 to -0,04668 | Yes | **** | <0,0001 |  |  |  |
| PM-S7a Rescue vs. Lyso-S7a Rescue | -0.02766 | -0,04414 to -0,01119 | Yes | **** | <0,0001 |  |  |  |
| PM-S7a Rescue vs. SV-S7a Rescue | -0.05508 | -0,07130 to -0,03886 | Yes | **** | <0,0001 |  |  |  |
| Lyso-S7a Rescue vs. SV-S7a Rescue | -0.02741 | -0,04362 to -0,01121 | Yes | **** | <0,0001 |  |  |  |
| Test details | Predicted (LS) mean 1 | Predicted (LS) mean 2 | Predicted (LS) mean diff, | SE of diff, | N1 | N2 | t | DF |
| WT vs. S7KO (+CRE) | 0.8669 | 0.8985 | -0.03159 | 0.005833 | 750 | 650 | 5.416 | 4148 |
| WT vs. S7a Rescue | 0.8669 | 0.8596 | 0.007326 | 0.005621 | 750 | 750 | 1.303 | 4148 |
| WT vs. PM-S7a Rescue | 0.8669 | 0.8674 | -0.0004883 | 0.005625 | 750 | 748 | 0.08681 | 4148 |
| WT vs. Lyso-S7a Rescue | 0.8669 | 0.895 | -0.02815 | 0.005621 | 750 | 750 | 5.009 | 4148 |
| WT vs. SV-S7a Rescue | 0.8669 | 0.9224 | -0.05557 | 0.005532 | 750 | 800 | 10.04 | 4148 |
| S7KO (+CRE) vs. S7a Rescue | 0.8985 | 0.8596 | 0.03892 | 0.005833 | 650 | 750 | 6.672 | 4148 |
| S7KO (+CRE) vs. PM-S7a Rescue | 0.8985 | 0.8674 | 0.0311 | 0.005837 | 650 | 748 | 5.329 | 4148 |
| S7KO (+CRE) vs. Lyso-S7a Rescue | 0.8985 | 0.895 | 0.003441 | 0.005833 | 650 | 750 | 0.5899 | 4148 |
| S7KO (+CRE) vs. SV-S7a Rescue | 0.8985 | 0.9224 | -0.02397 | 0.005748 | 650 | 800 | 4.171 | 4148 |
| S7a Rescue vs. PM-S7a Rescue | 0.8596 | 0.8674 | -0.007814 | 0.005625 | 750 | 748 | 1.389 | 4148 |
| S7a Rescue vs. Lyso-S7a Rescue | 0.8596 | 0.895 | -0.03548 | 0.005621 | 750 | 750 | 6.312 | 4148 |
| S7a Rescue vs. SV-S7a Rescue | 0.8596 | 0.9224 | -0.06289 | 0.005532 | 750 | 800 | 11.37 | 4148 |
| PM-S7a Rescue vs. Lyso-S7a Rescue | 0.8674 | 0.895 | -0.02766 | 0.005625 | 748 | 750 | 4.918 | 4148 |
| PM-S7a Rescue vs. SV-S7a Rescue | 0.8674 | 0.9224 | -0.05508 | 0.005536 | 748 | 800 | 9.948 | 4148 |
| Lyso-S7a Rescue vs. SV-S7a Rescue | 0.895 | 0.9224 | -0.02741 | 0.005532 | 750 | 800 | 4.955 | 4148 |

